# Single cell transcriptomic analysis reveals cellular diversity of murine esophageal epithelium and age-associated mitochondrial dysfunction

**DOI:** 10.1101/2021.02.08.430312

**Authors:** Mohammad Faujul Kabir, Adam Karami, Ricardo Cruz-Acuna, Alena Klochkova, Reshu Saxena, Anbin Mu, Mary Grace Murray, Kelsey Keith, Jozef Madzo, Hugh Huang, Jaroslav Jelinek, Tatiana Karakasheva, Kathryn E. Hamilton, Amanda B. Muir, Marie-Pier Tetreault, Kelly A. Whelan

## Abstract

Stratified squamous epithelium of the esophagus is comprised of basal keratinocytes that execute a terminal differentiation program in overlying suprabasal and superficial cell layers. Although morphologic progression coupled with expression of specific molecular markers has been characterized along the esophageal epithelial differentiation gradient, the molecular heterogeneity within the cell types along this trajectory has yet to be classified at the level of single cell resolution. To explore the molecular characteristics of esophageal keratinocytes along the squamous differentiation continuum, we performed single cell RNA-Sequencing transcriptomic profiling of 7,972 cells from murine esophageal epithelial sheets. We identified 8 distinct cell clusters in esophageal epithelium, unveiling an unexpected level of diversity, particularly among basal cells. We further mapped the cellular pathways and lineage trajectories within basal, suprabasal, and superficial clusters as well as within the heterogeneous basal cell populations, providing a comprehensive molecular view of esophageal epithelial cells in the context of squamous differentiation. Finally, we explored the impact of tissue aging upon esophageal epithelial cell clusters and demonstrated that mitochondrial dysfunction is a feature of aging in normal esophageal epithelium. These studies provide an unparalleled molecular perspective on murine esophageal keratinocytes that will serve as a valuable resource for dissecting cell type-specific roles in esophageal biology under conditions of homeostasis, aging, and tissue pathology.

## INTRODUCTION

In stratified squamous epithelium of the esophagus, basal cells give rise to overlying keratinocytes that exhibit a gradient of squamous differentiation as they move toward the lumen and ultimately desquamate. Squamous differentiation in esophageal keratinocytes is marked by downregulation of basal cell markers, including cytokeratins KRT5 and KRT14 (1, 2) and transcription factors SOX2 (3) and p63 (4), concomitant with induction of KRT13, KRT4, and Involucrin in early differentiation (suprabasal cells)(5, 6), then Filaggrin and Loricrin in late differentiation (superficial cells)(7, 8). Squamous differentiation is coupled to the cell cycle in the esophagus with proliferation restricted to basal cells (9). Presently, controversy exists regarding to what degree, if any, heterogeneity exists within basal esophageal keratinocytes (10-17). Maintenance of the exquisite differentiation gradient of esophageal epithelium is necessary for barrier function with its disruption being a hallmark of esophageal pathologies.

Age represents a well-established risk factor for development of esophageal lesions, both premalignant and malignant. Recent studies further demonstrate age-associated remodeling of esophageal epithelium via expansion of clones with mutations in cancer driver genes, including *NOTCH1* (18, 19). These genetic events become highly prevalent among physiologically normal human esophageal epithelium with age, despite a lack of gross alterations in tissue histology (18, 19). Although these studies provide valuable insight into the impact of tissue aging upon the mutational spectrum of esophageal epithelial cells, how aging influences the cellular landscape of epithelium at the transcriptional level has yet to be elucidated.

Single cell RNA-sequencing (scRNA-Seq) represents a powerful tool for unbiased, high-throughput interrogation of the distinct cell states associated with lineage commitment along the proliferation/differentiation axis in esophageal epithelium. In this study, we implemented this technique to provide the first comprehensive survey of murine esophageal epithelium in young and aged mice, revealing 8 transcriptionally distinct epithelial cell clusters: 5 basal, 1 suprabasal, and 2 superficial. We further performed in-depth characterization of the molecular features associated with both the 3 stages of lineage commitment in esophageal keratinocytes and the individual cell clusters comprising the basal, suprabasal and superficial compartments in murine esophageal epithelium. Pseudotemporal projection in the 8 epithelial cell clusters as well as in the 5 basal cell clusters provides an unprecedented level of resolution in esophageal epithelial cells as they traverse the basal-suprabasal-superficial continuum during squamous differentiation. We finally explored the impact of tissue aging upon the representation and transcriptional profiles of the 8 identified murine esophageal epithelial cell clusters, demonstrating for the first time that mitochondrial dysfunction is a feature of aged esophageal epithelium.

## RESULTS

### Molecular characterization of cell clusters along the squamous differentiation gradient in murine esophageal epithelium

To investigate the molecular heterogeneity of the esophageal epithelium at the level of single cell resolution, scRNA-Seq was performed using peeled esophageal epithelium from young (≤4 months) and aged (≥18 months) mice (**Figure 1A**). To maximize input data for initial studies, data from young and aged mice were integrated for dimensionality reduction. This minimized age-based effects and ensured that similar cell types across the age groups were grouped in the same clusters. Additionally, integration enabled direct comparisons of the representation and differential gene expression within clusters in young and aged mice. Seurat’s unsupervised dimensionality reduction and clustering workflow with Uniform Manifold Approximation and Projection (UMAP) of the 7,972 analyzed cells revealed 10 cell clusters with distinct transcriptional profiles (**Figure 1B, C**; **Supplementary Figure S1**). As our primary objective was to specifically evaluate esophageal epithelial heterogeneity, the 2 clusters displaying respective gene expression patterns consistent with fibroblasts or immune cells (**Supplementary Figure S2**) were excluded from downstream analyses. To reveal sources of heterogeneity within the data, the calculated UMAP dimensionality reduction was projected onto the entire epithelial dataset and the loadings were determined for all genes. *Krtdap* (encoding Keratin differentiation-associated protein) and *Krt5* (encoding Cytokeratin 5) were the genes that most significantly impacted cluster identification (**Figure 1D**). Based upon the continuum of highly *Krtdap*-positive cells to highly *Krt5*-positive cells across our epithelial dataset (**Figure 1E-G**) and the knowledge that KRTDAP and KRT5 have been established as respective markers of differentiated and basal cells in stratified squamous epithelium (2, 20), we utilized these markers to identify basal, suprabasal, and superficial cells within our dataset (**Figure 1C, D**).

**Figure 1.**
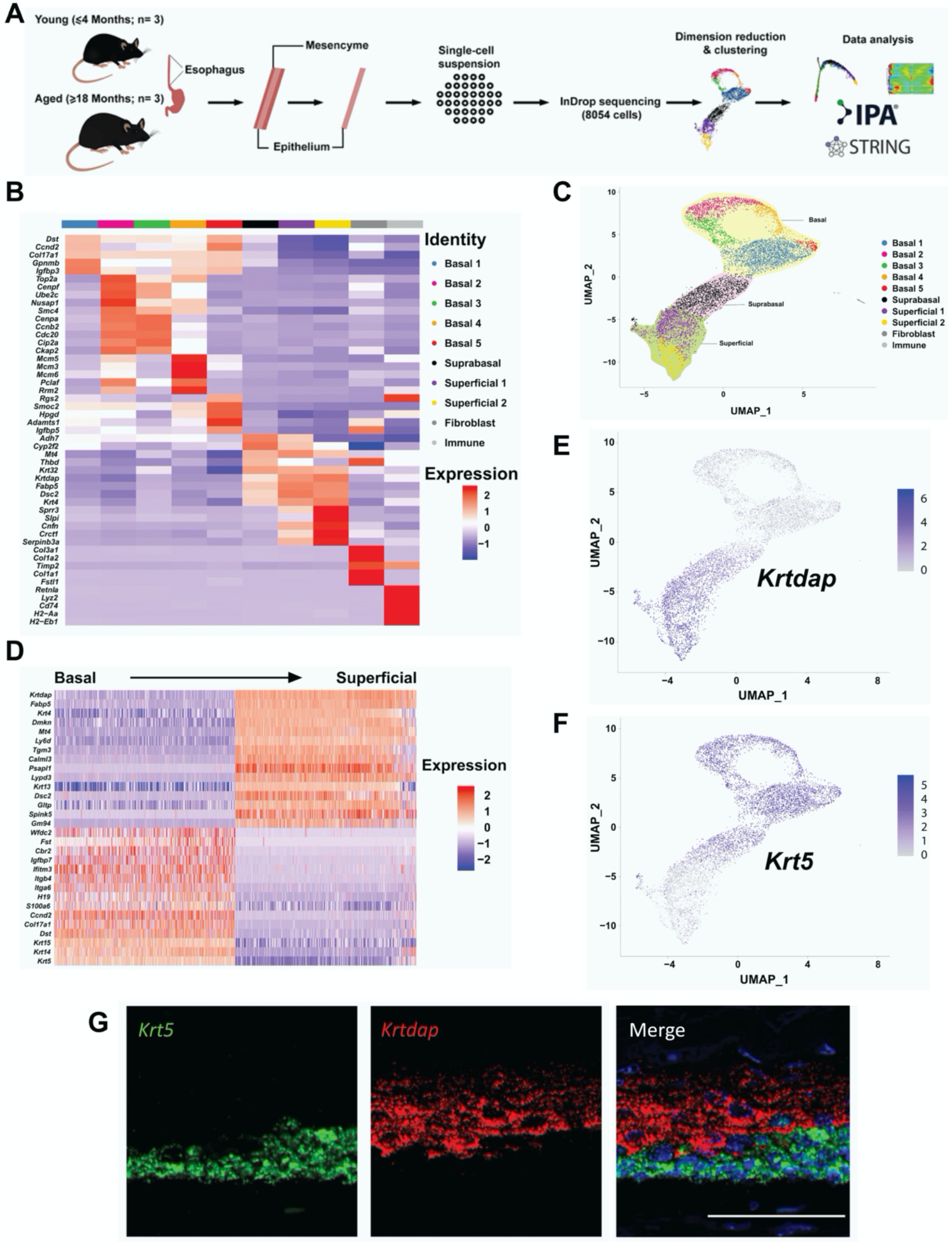
Identification of cell clusters in murine esophageal epithelium. (**A**) Schematic overview of experimental design. (**B**) Expression z-scores for the top 5 upregulated genes in each cluster. (**C**) Seurat’s Uniform Manifold Approximation and Projection (UMAP) was used to identify distinct cell clusters within the epithelial dataset. Eight epithelial communities were identified, as well as fibroblasts and immune cells. (**D**) Genes identified as primary contributors to UMAP based on their loadings are listed and their expression z-scores in cells across the epithelial dataset are shown. (**E, F**) Log1p normalized expression of the basal marker *Krt5* (**F**) and the superficial marker *Krtdap* (**E**) across the epithelial dataset is shown. (**G**) Representative image of RNA FISH to visualize *Krt5* and *Krtdap* in murine esophageal epithelium *in situ* (n=3 mice).

Examination of known molecular features associated with basal and differentiated esophageal keratinocytes supported our population classifications. As described with *Krt5*, the putative basal cell markers *Krt14, Krt15, Sox2* and *Trp63* revealed marked differential expression when comparing esophageal epithelial subsets identified as basal cells to those defined as suprabasal or superficial (**Figure 2A**). Additionally, *Mki67* expression was noted only in basal clusters 2, 3, and 4 (**Figure 2A**), consistent with restriction of proliferation to the basal cell compartment in esophageal epithelium. With regard to known markers of squamous differentiation, *Krt13* and *Krt4* were low in basal cell clusters with induction becoming apparent in superficial cells (**Figure 2A**). Expression of the late differentiation marker *Lor* (encoding Loricrin) was identified in 41% of cells in superficial cluster 2, with negligible expression found in all other clusters. Expression of *Flg* (encoding Filaggrin) and *Ivl* (encoding Involucrin) was largely undetectable across all clusters in our dataset (**Figure 2A**). Unbiased determination of transcripts displaying specificity at each stage of lineage commitment was also performed (**Supplementary Figure S3**), identifying both known and novel markers for basal, suprabasal and superficial subsets. Immunostaining provided validation of COL17A1, ATP1B3, and CNFN as novel markers of basal, suprabasal and superficial cells, respectively, in murine esophageal epithelium (**Figure 2B**). These novel markers of the 3 stages of esophageal lineage commitment also displayed appropriate localization along the basal/differentiated cell axis in a scRNA-Seq dataset performed on human esophageal tissue specimens (**Supplementary Figure S4**)(21). Gene Ontology (GO) analysis of differentially expressed genes (DEGs) in basal, suprabasal, and superficial cell clusters further provided insight into the dynamic molecular signatures associated with lineage commitment in esophageal keratinocytes (**Supplementary Figure S5A-D**).

**Figure 2.**
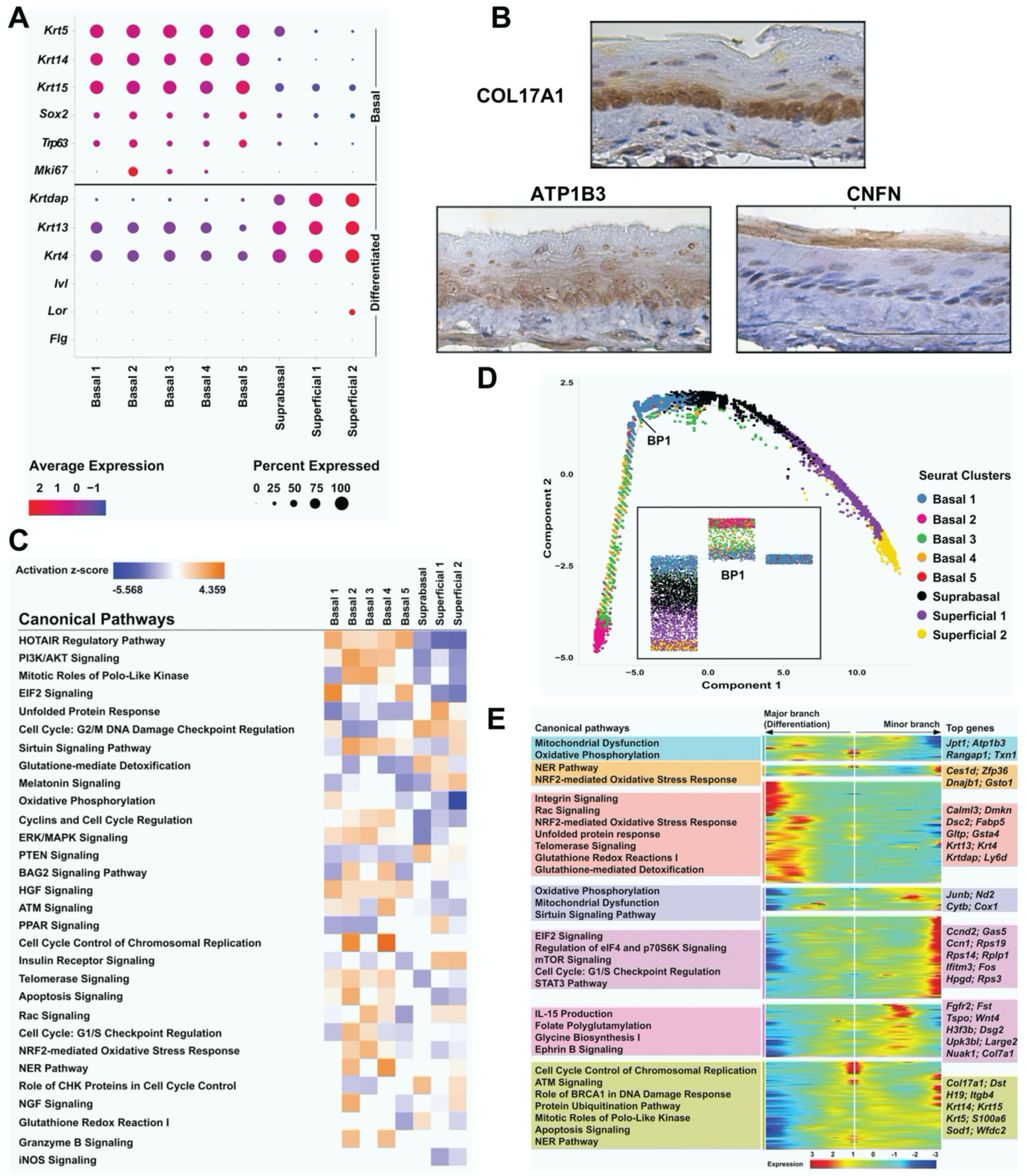
Molecular characterization of the basal/differentiated cell axis in murine esophageal epithelium. (**A**) Cluster-average expression z-scores of putative basal and differentiated markers. Circle size reflects percentage of cells with non-zero expression level for indicated genes. Color intensity reflects average expression level across all cells within each cluster. (**B**) Immunohistochemistry for indicated proteins in murine esophageal epithelium. (**C**) IPA prediction of canonical pathways and their activation state in each cluster. In heat map, ranking is based upon activation z-score and color intensity depicts relative value of activation z-score. (**D**) Pseudotime trajectory derived from Monocle’s reversed graph embedding. Each cell is colored according to cluster. Inset displays minimum spanning tree structure of the basal-superficial axis pseudotime. BP1, branchpoint 1. (**E**) Branch-dependent analysis of branch point 1. The heatmap represents the expression z-scores of genes along the trajectory. IPA revealed pathways significantly altered in gene clusters. Representative pathways are shown.

We next aimed to more comprehensively characterize the molecular features associated with individual cell clusters in esophageal epithelium. Although Ingenuity Pathway Analysis (IPA) of the DEGs in each cluster revealed overlap, cluster-specific pathway signatures were also detected lending further support to the individual molecular identity of the 8 identified clusters (**Figure 2C, Supplementary Figure S6**). To gain insight into the relationships existing between individual esophageal epithelial cell clusters along the proliferation/differentiation axis, we next employed pseudotemporal trajectory inference mapping of the clusters comprising our dataset. This analysis identified an organized progression from basal progenitor cells to suprabasal progeny and ultimately terminally differentiated esophageal keratinocytes, which was confirmed by expression mapping of putative basal and differentiated markers along the predicted trajectory (**Figure 2D; Supplementary Figure S7A, B**). The pseudotemporal projection was rooted in basal cluster 2, which displays both the most robust expression of *Mki67* and suppression of differentiation markers (**Supplementary Figure S7B, C**), followed by mixed representation of basal clusters 3 and 4, then an enrichment for basal cluster 1 (**Figure 2D**). A branch point was identified in the area of basal cluster 1 enrichment, with a minor branch consisting of basal clusters 1 and 5 diverging from the primary branch that leads toward terminal differentiation (**Figure 2D**). Notably, the primary trajectory continued to progress in a linear fashion, with suprabasal cells giving rise to superficial cluster 1, which ultimately led to superficial cluster 2 (**Figure 2D; Supplementary Figure S7B**). Branch expression analysis modeling (BEAM) revealed distinct gene expression profiles in the two branches stemming from branch point 1 with differentiation-associated genes, including *Krtdap, Krt13* and *Krt4*, displaying upregulation in the primary branch leading toward terminal differentiation (**Figure 2E**). Additionally, IPA provided insight into the pathways that are altered in the primary and minor branches (**Figure 2E**). In sum, these studies unveil previously unappreciated cellular heterogeneity within murine esophageal epithelium and identify potential regulators of the establishment and maintenance of this heterogeneity at the levels of both genes and pathways.

### Defining transcriptional heterogeneity and mapping cell fate trajectories in esophageal basal cells

Given the marked level of molecular heterogeneity found within basal cells, we continued to perform pseudotemporal trajectory analysis specifically using the 5 basal cell clusters of murine esophageal epithelium. Upon removal of suprabasal and superficial clusters, a pseudotemporal projection with multiple branches was identified (**Figure 3A**). Of the 4 branch points arising from the prebranch, 3 gave rise to minor terminal branches (**Figure 3A**). BEAM identified gene expression profiles for each of these branches with the branch emerging from branch point 1 displaying enrichment of cell cycle-associated genes and the branches emerging from branch points 2 and 3 displaying enrichment of genes associated with cell differentiation and development (**Supplementary Figure S8A-C**).

**Figure 3.**
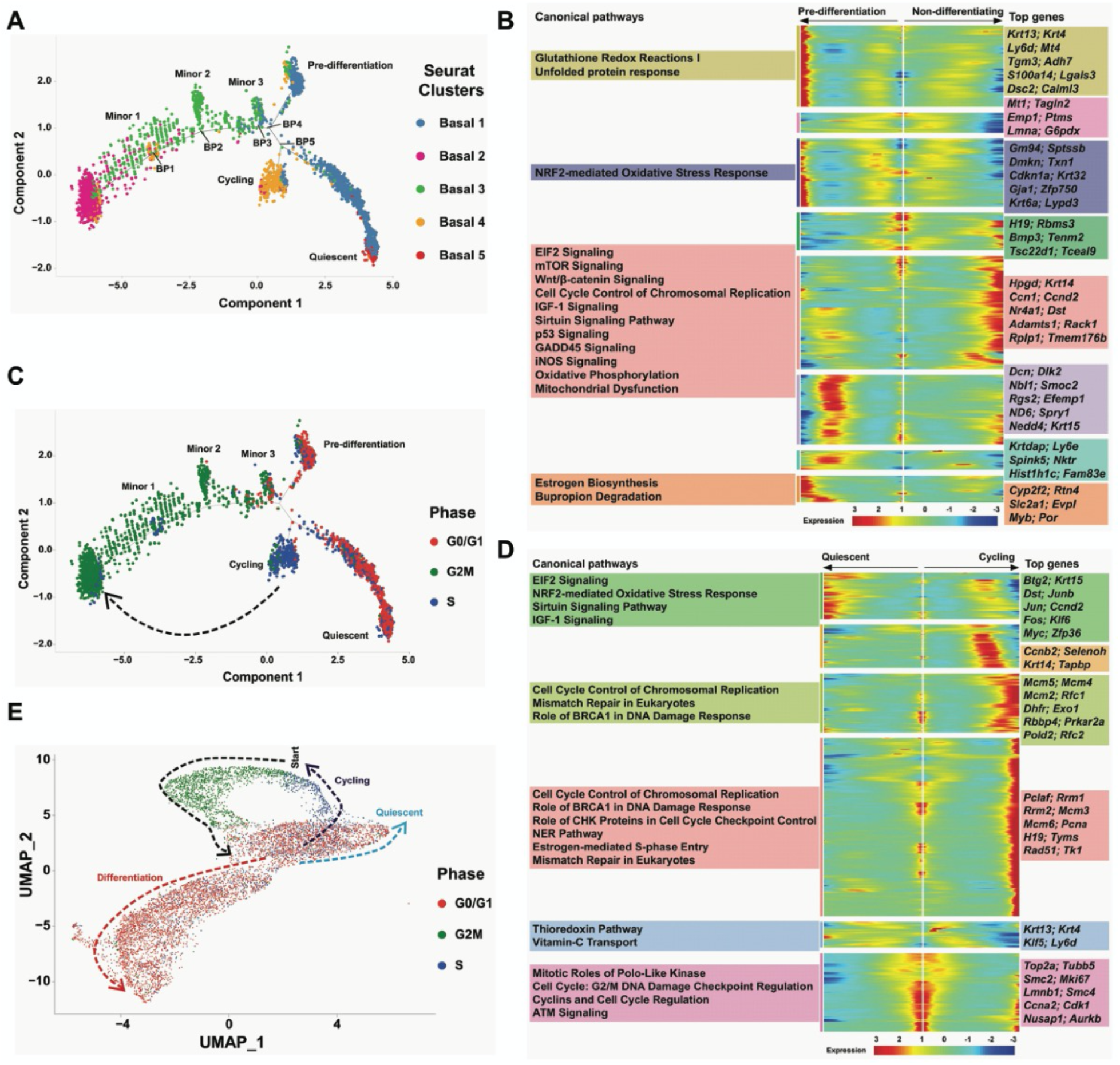
Inference of relationships between basal cell subtypes in murine esophageal epithelium by pseudotime. (**A**) Pseudotime projection of basal epithelial cell clusters as previously identified in Seurat. BP, branch point. (**B**) Branch Expression Analysis Modeling (BEAM) at branch point 3. Heatmap represents expression z-scores of genes along the basal cell trajectories defined as pre-differentiation or non-differentiating. Ingenuity Pathway Analysis (IPA) revealed pathways significantly altered in gene clusters with representative pathways shown. (**C**) Cell cycle phases of each cell in the pseudotime projection of the basal epithelium. The cell cycle phases were determined based on the upregulation S and G2M marker genes. Cells with lower expression of S and G2M markers were categorized as G0/G1 phase cells. (**D**) BEAM of branch point 5. Heatmap represents expression z-scores of genes along the basal cell trajectories defined as cycling or quiescent. IPA revealed pathways significantly altered in gene clusters with representative pathways shown. (**E**) Calculation and comparison of cell cycle phase markers reveals the cell cycle phases of each cell in the UMAP projection of the esophageal epithelium. Start indicates location of basal cluster 2, the initiation point for both pan-epithelial and basal pseudotemporal projections, which then proceeds from G2/M phase to G0/G1 phase as indicated by the black arrow. Colored arrows represent predicted paths for listed indicated fates in esophageal epithelium

The first primary branch of the pseudotemporal analysis emerged at branch point 4 (**Figure 3A**). We termed this branch ‘Pre-differentiation’ given its marked transcriptional similarity to basal cells in the branch giving rise to terminally differentiated keratinocytes in our pan-epithelial pseudotemporal analysis (**Supplementary Figure S9**). Upregulation of differentiation-associated genes, including *Krt4, Krt6a, Krtdap, Ly6d, Dmkn* and *Gja1*, was also noted in the pre-differentiation branch (**Figure 3B**). IPA further demonstrated enrichment for glutathione redox reaction I, unfolded protein response (UPR) and nuclear factor erythroid 2–related factor 2 (NRF2)-mediated oxidative stress response pathways (**Figure 3B**), all of which were also enriched in differentiating keratinocytes (**Figure 2E**). These findings support a model wherein cells comprising the pre-differentiation branch represent basal cells that are poised to undergo squamous cell differentiation.

The second branch emerging from branch point 4, which we collectively termed ‘Non-differentiating’, was characterized by signaling related to proliferation, Wnt, insulin growth factor (IGF)-1 and sirtuins, as well as mitochondrial dysfunction and oxidative phosphorylation (**Figure 3B**). This branch subsequently gave rise to the 2 remaining major branches in the esophageal basal cell pseudotemporal trajectory (**Figure 3A**). We termed one of these branches as ‘Cycling’ based upon its expression of genes associated with proliferation, including *Mcm2*/*3*/*4*/*5*/*6, Rrm1*/*2, Rfc1*/*2* and *Pold2*, and enrichment of pathways related to cell cycle control and DNA repair (**Figure 3D**). Consistent with these data, cell cycle analysis in the basal pseudotime revealed an enrichment of S phase genes in the cycling branch (**Figure 3C**). As pseudotime fails to account for cyclical processes, we further evaluated the cell cycle in the context of the original cell cluster map (**Figure 3E**). The pan-epithelial and basal pseudotemporal analyses initiate in basal cluster 2 which is enriched for G2/M genes (**Figure 3E**). Cells in basal cluster 3 remain in the G2/M phase of the cell cycle before giving rise to G0/G1-enriched basal cluster 1 (**Figure 3E**). Basal cluster 1 cells may then remain in G0/G1 phase as they execute the terminal differentiation program or they may re-enter the cell cycle, progressing to the S phase (**Figure 3E**). Thus, cell cycle analyses in **Figure 3D** and **3E**, suggest that the cycling population identified in the basal pseudotime may not represent a terminal cell fate, but rather may be composed of cells that will progress into G2/M as part of the prebranch (**Figure 3D, E**). Notably, despite the association between cell cycle gene expression and esophageal cell identity, regression of cell cycle genes from the dataset minimally impacted cluster identity (**Supplementary Figure S10**).

The final branch in the basal cell pseudotime displayed enrichment for markers of the G0/G1 phase of the cell cycle (**Figure 3C**). An inhibition of cell proliferation in this branch was supported by downregulation of markers associated with proliferation (*Pcna, Mki67)* and DNA replication (*Rad51, Mcm2/3/4/5*), coupled with upregulation of proliferation inhibitors (*Btg2, Klf6, Gas1)* (**Figure 3D; Supplementary Figure S11**). We termed this branch ‘Quiescent’ based upon these data in conjunction with upregulation of genes that have been associated with quiescence in various experimental systems, including *Junb, Zfp36l1, Btg2, Spry1, Gas1, Dcn*, and *Nr4a1* (22-29) (**Figure 3C, D; Supplementary Figure S11**), and low RNA content (**Supplementary Figure S12**). IPA further revealed enrichment for signaling associated with eukaryotic initiation factor 2, NRF2, sirtuins and IGF-1 in the quiescent branch of the basal pseudotime (**Figure 3D**). Notably, the quiescent branch of the basal cell pseudotime displayed remarkable transcriptional similarity to the minor branch in the pan-epithelial pseudotime (**Supplementary Figure S13**), indicating that esophageal epithelial cells have 2 potential terminal cell fates: differentiation or quiescence.

### Defining age-associated alterations in esophageal epithelial biology

We finally aimed to define the impact of tissue aging upon the cellular and molecular landscape of esophageal epithelium. Relative cluster representation was impacted by age with significant enrichment of basal cluster 5 and suppression of basal clusters 3 and 4 found in aged mice as compared to their young counterparts (**Figure 4A, B**). These data were of particular interest as basal cluster 5 is enriched at the terminal end of the quiescent branch of the basal pseudotime while basal cluster 4 is the predominant cell type found in the cycling branch of the predicted basal trajectory (**Figure 3A, C**). We continued to utilize 3D esophageal organoid culture to determine if age may functionally impact basal cell behavior. Organoid formation rate upon initial plating was similar in esophageal epithelial cells isolated from young and aged animals (**Figure 4C**). Upon passaging, however, organoid formation capacity increased in cultures from aged mice while decreasing in those from young mice (**Figure 4C**), supporting altered basal cell dynamics as a feature of esophageal aging.

**Figure 4.**
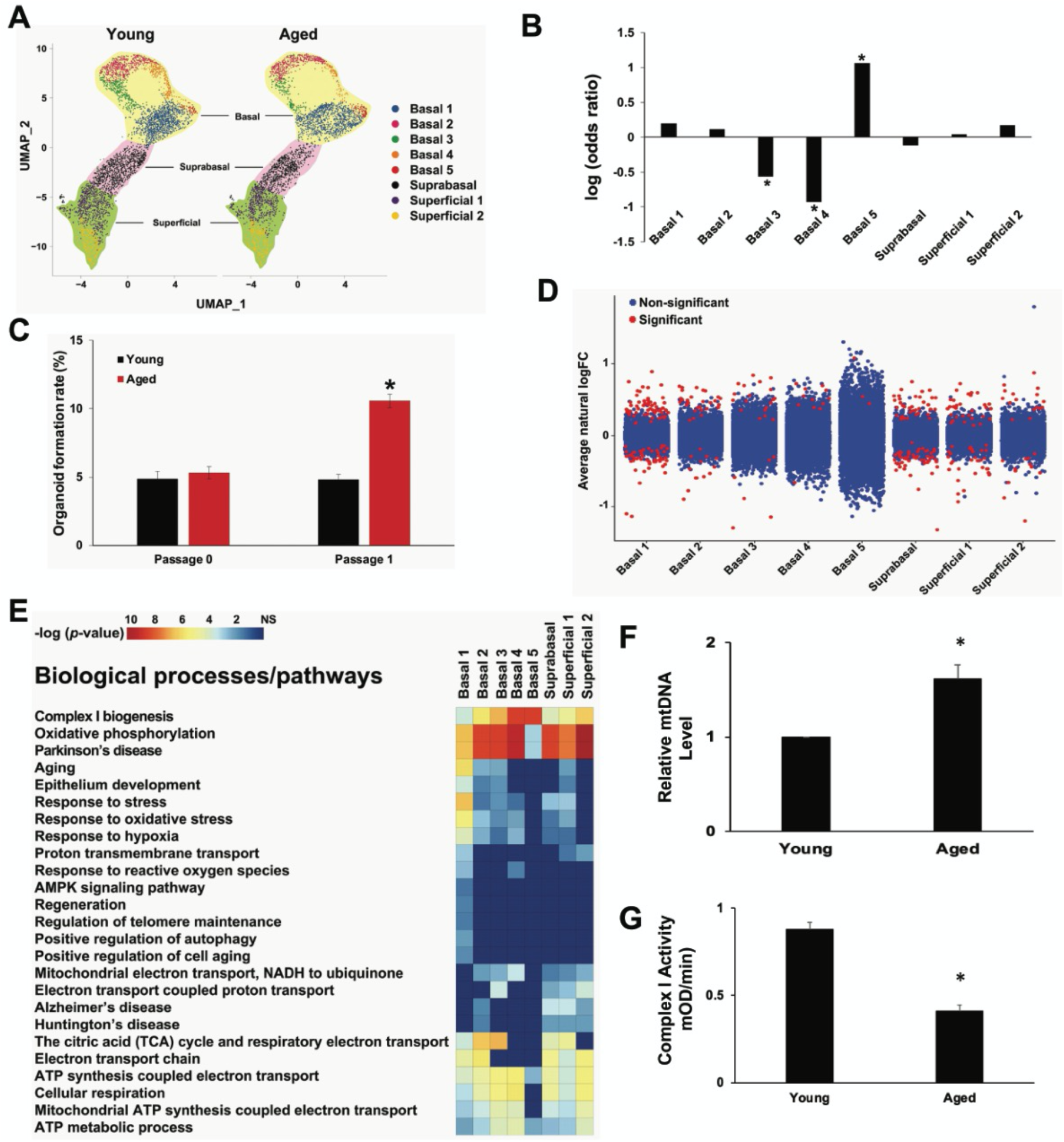
Age-associated alterations in murine esophageal epithelium. (**A**) UMAP visualization of Seurat cell clusters in young and aged esophageal epithelium. (**B**) Proportion of each cell cluster as a fraction of all cells in the epithelium for each age group. Log_2_ of the odds ratio of fractional proportions of each cluster. Odds ratio calculated by fractions in each respective cluster in aged mice over young mice. (**C**) Epithelial cells isolated from esophagi of young and aged mice were subjected to 3D organoid culture (passage 0). After 14, days organoids were dissociated and re-seeded for 3D organoid culture (passage 1). Organoid formation rate (number organoids formed/number cells plated) was determined after 14 days of growth. *, p<0.0001. (**D**) Average natural log fold change (FC) of each gene in each epithelial cluster, aged cells compared to young cells in each respective cluster. (**E**) STRING analysis identified the biological processes and pathways significantly altered in esophageal epithelial clusters in aged mice relative to young mice. (**F**) Expression of mitochondria-encoded ND1 gene was determined relative to nuclear-encoded gene Ikbβ in peeled esophageal epithelium from young and aged mice. *, p<0.005. (**G**) Mitochondrial complex I activity was measured in esophageal tissue lysates from young and aged mice.*, p<0.0001.

We further evaluated the impact of aging upon gene expression within epithelial cell clusters. Although cell identity was largely preserved in aged esophageal epithelium, cluster-specific alterations were identified to a varying degree (**Figure 4D; Supplementary Figure S14A**). Enrichment of genes associated with mitochondrial electron transport chain (ETC) complex I biogenesis and oxidative phosphorylation was identified across esophageal epithelium (**Figure 4E**). Indeed, of the 64 genes displaying age-associated alterations in gene expression, 11 were associated with mitochondrial biology (**Supplementary Figure S14B**). Age-associated disruption of mitochondrial biology was confirmed as esophageal epithelium of aged mice displayed increased mitochondrial DNA content as well impaired mitochondrial complex I activity (**Figure 4F, G**).

## DISCUSSION

While recent advances in scRNA-Seq technology have facilitated molecular characterization of the cell types comprising a wide variety of tissues, the current study represents the first in-depth investigation of normal esophageal epithelium at the single cell level. This high throughput scRNA-Seq analysis reveals marked cellular diversity in murine esophageal epithelium with the heterogeneity of individual cell clusters apparent at the level of biological processes. Basal cluster 1, for example, uniquely displayed activation of oxidative phosphorylation. Among all clusters, basal cluster 2 was characterized by the most significant activation of signaling associated with nerve growth factor, which of particular interest as Neurotrophin receptor p75 has been linked to esophageal stem cells (30). HOTAIR regulatory pathway, which was activated across basal clusters and inhibited in suprabasal and differentiated clusters, exhibited the greatest enrichment in basal cluster 5, and was also predicted to be inhibited in both superficial populations. With regard to the suprabasal cluster, inhibition of PI3K/Akt signaling coupled with activation of PTEN further suggested suppression of the PI3K pathway. Additionally, activation of glutathione-mediated detoxification in suprabasal cells indicated potential alterations in cellular antioxidant responses. Differential alterations in several pathways were also noted when comparing the 2 superficial clusters. Peroxisome proliferator-activated receptor signaling and UPR were activated in superficial 1 where inducible nitric oxide synthase and hepatocyte growth factor (HGF) signaling were inhibited, while superficial cluster 2 displayed unique activation of melatonin signaling. These data support the individual molecular identity of the 8 identified subsets of esophageal keratinocytes and additionally present HGF, sirtuins, HOTAIR, all of which have been implicated in esophageal cancer (31-36), as pathways of interest with regard to regulation of esophageal homeostasis.

The presence of 5 basal cell subsets in murine esophageal epithelium of is of particular interest as the question of what degree, if any, heterogeneity exists among esophageal basal cells remains controversial. Although studies using lineage tracing in mice coupled with mathematical modeling support a single-progenitor model wherein all esophageal basal cells have equal capacity to proliferate or differentiate (13, 17), several other studies have identified markers associated with functional heterogeneity in the mouse esophagus. Slow-cycling/long-lived esophageal basal cells with self-renewal capacity have been identified by positivity for CD34, KRT15, or a combination of high integrin a6 and low CD71 expression (11, 14, 15). By contrast, work by DeWard et al. indicates that a proliferative subset of basal cells defined by high expression of integrin β4 and positivity for CD73 exhibits the greatest stem cell potential in murine esophageal epithelium (12). Employing pseudotemporal analysis, we have developed a model wherein G0/G1-enriched basal cluster 1 cells diverge to give rise one of 3 potential cell fates: differentiation, proliferation/renewal or quiescence. Based upon our analysis of the transcriptional programs of the first 2 primary branches emerging in the basal pseudotemporal trajectory, we propose that G0/G1 basal cells either commit to execution of the squamous differentiation program or are presented with a subsequent decision point at which cells that remain in G0/G1 enter into a quiescent state, while cells that progress to S phase serve as a reservoir for repopulation of the basal 2-rich progenitor pool. We further postulate that the minor branches stemming from the pre-branch may represent basal cells that fail to effectively execute the delineated cell fate determinations. Our findings are not inconsistent with published data on esophageal basal cell heterogeneity as we note the presence of both a proliferative progenitor pool and a population with features of quiescence. Additionally, as it been demonstrated that while esophageal epithelial cells exhibit equipotent potential to proliferate or differentiate in order to facilitate tissue renewal, these cells may transiently shift toward a more proliferative state during tissue repair (13). Future studies will investigate the impact of tissue-damaging agents on the molecular landscape of murine esophageal epithelium and also define specific markers of the identified cell populations in order to explore functional roles for these populations under conditions of homeostasis, damage, and disease by employing lineage tracing along with ablation strategies.

Our findings further indicate that tissue aging has an impact on esophageal basal cell dynamics as well as mitochondrial biology. We report that basal clusters 3 and 4 are significantly underrepresented in aged mice while basal cluster 5 is overrepresented. These data support an age-associated decrease in proliferation as basal clusters 3 and 4 show respective enrichment for G2/M and S phase genes, and basal 5 exhibits G0/G1 gene enrichment as well as localization at the tip of the quiescent branch of the basal pseudotime projection. Organoid assays demonstrated increased passaging capacity in esophageal epithelium of aged mice, supporting altered basal cell dynamics with age, potentially via enrichment of slow-cycling cells with enhanced self-renewal capability. Esophageal cancer incidence increases with age and peaks in the seventh and eighth decades of life (37). Additionally, it been demonstrated that age-associated remodeling of physiologically normal esophageal epithelium occurs via expansion of clones with mutations in cancer driver genes (18, 19). As such, it is tempting to speculate that basal cluster 5 may represent a subpopulation of basal cells that is capable of acquiring the mutational spectrum necessary for initiating esophageal carcinogenesis owing to its long-lived nature. Finally, across esophageal epithelium we found enrichment of genes associated with mitochondrial ETC complex I biogenesis and oxidative phosphorylation in aged mice with functional analysis of mtDNA demonstrating an increase. Complex I activity, however, was found to be decreased in aged esophageal epithelium, potentially as gene enrichment represents an attempt to restore complex I function. Mitochondrial complex I is the initial point of entry for electrons into the ETC, contains the cellular NAD+ pool, and alterations in its function are a contributing factor in various diseases, including cancer (38). Defects in complex I have been associated with alterations in redox homeostasis (39), raising the possibility that decline in complex I activity may facilitate age-associated accumulation of mutations in esophageal epithelium. Presently, the role of mitochondria in esophageal epithelial homeostasis remains elusive and should be further explored.

## MATERIALS & METHODS

### Murine Epithelial Tissue Collection and Processing

Wild type C57BL/6 mice (Cat# 000664) were purchased from Jackson Laboratories at age 12-14 weeks or 72 weeks. Mice were allowed to acclimate for at least 2 weeks prior to use for experiments. Whole esophagi were dissected from young (≤3 months; Range 14-16 weeks) and aged (≥18 months; Range 18-20 months) mice. For experiments using peeled esophageal epithelium, epithelium was physically separated from underlying submucosa using forceps then the esophagus was cut open longitudinally to expose the epithelial surface. For single cell isolation, peeled esophageal epithelium of 3 young (16 weeks of age) and 3 aged (72 weeks of age) male mice was incubated in 1 ml of 1X Dispase I in Hank’s Balanced Salt Solution (HBSS) for 10 minutes at 37° C with shaking at 1,000 RPM. Following removal from Dispase I, esophageal epithelium was chopped into 3 pieces with sharp scissors then incubated in 1 ml of 0.25% Trypsin-EDTA for 10 minutes at 37° C with shaking at 1,000 RPM. Trypsin and tissue pieces were forced through a cell strainer (70 μm) into a 50 ml conical tube containing 4 ml of soybean trypsin inhibitor (STI). Cells were pelleted at 1,200 RPM for 5 minutes then resuspended in 500 μl of the complete mouse keratinocyte–serum-free medium without calcium chloride (Gibco Cat# 37010022). Cell number and viability were measured by Automated Cell Count (Invitrogen Countess II FL) by mixing 10 μl of cell suspension with 10 μl 0.4% trypan blue solution (1:1). For single-cell experiments, at least 300,000 cells with 85% viability were isolated from each mouse, serving as individual biological replicates. Cells were kept on ice until the time of single cell encapsulation. For downstream molecular studies, peeled esophageal epithelium or whole esophagus from equal numbers of male and female mice were processed as described below.

### scRNA Library Preparation and Sequencing

The inDrop™ System and single-cell RNA sequencing kit (1CellBio, Watertown MA) were used for single cell encapsulation. Approximately 2,000 cells were co-encapsulated in 1-3 nL droplets with barcode-labeled hydrogel beads and reverse transcription/cell lysis reagents following the manufacturer’s protocol. Oligonucleotide primers with cell barcodes were released from the beads by an UV light exposure and reverse transcription was performed in the emulsion of droplets in oil. Barcoded cDNA was released by breaking the droplets after reverse transcription and stored at -80°C. Library preparation was performed following the manufacturer’s instructions (1CellBio protocol version 2.3). Libraries were sequenced at Fox Chase Cancer Center Genomic Facility on an Illumina HiSeq 2500 instrument using paired-end 2 x 50 base pairs.

### Deconvolution of scRNA-Seq Reads

FASTQ files from the sequencing run were downloaded from Illumina’s BaseSpace sequence hub. To identify unique cellular barcodes in each sequencing run, UMI-tools (40) was used to whitelist and extract the barcodes. Likely barcodes were found using the whitelist function of UMI-tools, which searches for the inDrop regular expression pattern of “(?P<cell_1>.{8,12})(?P<discard_1>GAGTGATTGCTTGTGACGCCTT){s<=2}(?P<cell_2>.{8})(?P<umi_1>.{6})T(42).*” in the R2 of the read pairs (40). Using a whitelist of likely barcodes, the extract function relocated both the cell barcode and the unique molecular identifier found in the same read to the read name in the FASTQ files. The extraction retains the information of unique cellular barcodes while enabling correct read mapping of endogenous genes downstream without the added cell and transcript identifiers. Reads were then mapped using the aligner STAR v2.7 (41). First, the murine genome index was made using the M23 GRCm38 genomic sequence from GENCODE (https://www.gencodegenes.org/mouse/release_M23.html). Using the index, all read pairs were aligned to the genome for an output of BAM files. The resulting alignments were then mapped to the murine gene transfer file (GTF) from GENCODE using featureCounts to count the number of reads mapping to each gene in the genome. Finally, the UMI-tools count function was used to summarize the gene counts in each cell in each sample to give an output of a matrix with gene names and cell barcodes.

### Data Filtering, Integration, Dimensionality Reduction, and Clustering

The matrices for each sample were imported and transformed into objects for further processing (42). Genes expressed in 3 or fewer cells were excluded from analysis. To eliminate dead cells or doublets, cells with the expression of less than 250 unique genes or over 2,500, respectively, were excluded. Additionally, cells with over 20% of their transcripts consisting of mitochondrial genes were excluded. 7,972 cells remain after filtering, compared to 10,701 before. Analysis of the filtered matrices follows the Seurat integration workflow described by Stuart, *et. Al* (43). Each transcript count in a cell was ‘LogNormalized’ (divided by the total number of genes in that cell, and then multiplied by a scale factor of 10,000). The resulting number was then natural-log transformed with log1p. For each matrix, the top 2000 variable genes were calculated using variance-stabilizing transformation. CCA was then used to find integration transcript anchors between all of the matrices. Genes used for integration were ranked by the number of matrices they appear in. The matrices were then merged according to the integration genes and re-normalized using the LogNormalize procedure. From this point on, dimensionality reduction used the genes and values that were pre-processed using the integration workflow. However, the data unused for integration was retained for differential gene testing in later steps. The resulting dataset was then scaled and centered for dimensionality reduction. PCA was used for initial dimensionality reduction and later for clustering, resulting in 30 principal components. The PCA principal components were then used as input to the UMAP dimensionality reduction procedure (arXiv:1802.03426), using 30 neighbors for local neighborhood approximation and embedding into 2 components for visualization. Because of our interest in the relationship between cell cycle phases and cell fates, we opted not to regress cell cycle genes in our dimensionality reduction steps. A Shared Nearest Neighbor (SNN) graph was then constructed with the principal components of PCA by first determining the 20 nearest neighbors for each cell and subsequently creating the SNN guided by the neighborhood overlap between each cell and 20 of its nearest neighbors. Clusters were then determined by a modularity optimization algorithm by Waltman and van Eck (44).

### Cell Cluster Analyses

For each cluster, DEGs were calculated by comparing the expression of genes within the cells of the cluster over the expression of the genes in all other clusters. The statistical workflow to determine differential expression was Seurat’s implementation of Model-based Analysis of Single-cell Transcriptomics (MAST) by Finak, *et. a*l (45), a GLM-based framework that treats cellular detection as a covariate. The significance cutoff for DEGs is a Bonferroni-adjusted p-value of 0.05, and the fold change cutoff is below -0.25 or above 0.25 natural log fold change. To characterize the differential regulation of pathways in each cluster, DEGs that pass the cutoff from each cluster were exported into IPA (http://www.ingenuity.com) (Qiagen) for core analysis. In addition, GO biological processes significantly altered in esophageal clusters were categorized by PANTHER (46). For pseudotemporal inference, cells analyzed with Seurat were exported to Monocle (47) for analysis in a CellDataSet structure. Pre-processing for Monocle included estimation of size factors and dispersion and choosing genes to be used for ordering the cells in pseudotime and clustering according to the “dpFeature’’ procedure. Genes expressed in less than 5% of cells were excluded in the ordering and clustering set. To see if Monocle’s own dimensionality reduction method can corroborate Seurat’s UMAP based workflow, dimensionality reduction was performed in Monocle as well. t-distributed stochastic distributed neighbor embedding (tSNE) was done using log normalization on the scaled gene counts, and clusters were determined with density peak clustering with a k of 50. A differential gene test was done to distinguish genes that were significantly differentially expressed in different clusters using a model based on vector generalized linear and additive models (VGAM) in its R implementation (https://www.stat.auckland.ac.nz/~yee/VGAM/doc/glmgam.pdf). To order cells, 2000 genes with the lowest q-value from the differential gene test was chosen. The genes are then used for dimensionality reduction on the cells using the “DDRTree” algorithm (47), Monocle v2’s built-in reversed graph embedding workflow for pseudotemporal trajectory inference. The trajectory resolved a tree with three branches: a branch in which differentiation takes place, ending with cluster Superficial 2; a minor branch where Basal 5 is enriched; and a basal branch where the proliferation marker *Mki67* is enriched, ending with Basal 2. While all terminal endpoints of the main pseudotime trajectory can be set as the “root state” of cells, we determined the root state to be the proliferative branch based on canonical knowledge dictating that basal cells transition from proliferative to differentiated. Rooting at a point of proliferation was then also used in the basal-only pseudotime analysis. Monocle’s branch expression analysis modeling (BEAM) workflow was implemented in branch points of trajectories. BEAM performs a statistical test to see whether or not a gene’s mean expression changes smoothly as a function of pseudotime and if the mean expression’s curves are significantly different in each branch. The expression of genes that were identified as significantly branch-dependent (q-value of < 0.05) were observed and hierarchically clustered to identify groups of genes with similar expression changes across pseudotime. The canonical pathways and biological processes significantly altered in BEAM clusters were identified by IPA and STRING tools (48). To assess quiescence in pseudotime, total transcripts per cell and expression of genes upregulated in quiescence and stemness were analyzed. A gene list from Cheung and Rando (49) was used to score the expression of quiescence and stemness genes as a percentage of total expression in each cell. The Seurat function CellCycleScoring was used to predict the cell cycle of each cell. The function takes as input a list of S phase upregulated genes and G2/M upregulated genes, and outputs the score for each phase. The S and G2/M phase genes were provided within the Seurat package as objects “cc.genes.updated.2019$s.genes” and “cc.genes.updated.2019$g2m.genes”, respectively. The cell cycle phase is determined by the dominating score. Cells with weak scores for both phases are classified as G0/G1 phase cells. To compare each clusters’ proportional size between the different age groups, cells from young mice and aged mice in each cluster were identified and their proportional change as a fraction of total cells for each age group was tested for significance using a binomial test. The p-value for each cluster was then corrected with the Bonferroni procedure to account for multiple comparisons. STRING analysis was performed to identify the biological processes and pathways significantly altered with age in clusters or across the entire dataset (48). For validation of clusters, genes with the most cluster-exclusive expression were selected. For each cluster, genes were first ranked by their average expression in clusters other than the cluster of interest in ascending order and subsequently by fold change.

### In situ studies

RNA fluorescence in situ hybridization (FISH) and immunohistochemistry (IHC) were performed in formalin-fixed paraffin-embedded murine esophageal specimens. IHC was performed for COL17A1 (Invitrogen, MA5-24848l 1:100), ATP1B3 (Abcam, ab137055; 1:100) and CNFN (Novus Biologicals, NBP2-14668; 1:100) using a standard protocol as previously described (50). Slides were counterstained with Hematoxylin and imaged on a Leica DM30 microscope at 400X magnification. RNA FISH was performed using RNAscope technology (Advanced Cell Diagnostics) following the manufacturer’s protocol and RNAscope probes for murine *Krt5, Krtdap*, positive control, and negative control. Slides were counterstained with DAPI and imaged on a Leica SP8 confocal microscope at 400X magnification.

### 3D Organoid Assays

Murine esophageal 3D organoid formation assays were performed on freshly isolated primary murine epithelial cells (PMECs) as previously described (50). Briefly, a single cell suspension of PMECs in keratinocyte serum free medium was mixed with 90% Matrigel. For each well of a 24-well plate, 500 cells in 50μl Matrigel were seeded to initiate 3D organoid formation. After solidification, 500 μl of advanced DMEM/F12 supplemented with 1X Glutamax, 1X HEPES, 1X penicillin-streptomycin, 1X N2 Supplement, 1X B27 Supplement, 0.1 mM N-acetyl-L-cysteine, 50 ng/ml human recombinant EGF, and 2.0% Noggin/R-Spondin-conditioned media was added and replenished every 3-5 days. At the time of plating, 10 μM Y27632 was added to the culture medium. Organoids grown for 14 days were subjected to phase-contrast microscopy and counting using a Lionheart FX automated microscope (BioTek Instruments, Inc.). Organoid formation rate was determined as follows [(number of organoids at day 14/number of cells plated at day 0) x 100]. For passaging, organoids were recovered from Matrigel following incubation for 5 minutes with Dispase I then incubated in 1 ml of 0.25% Trypsin-EDTA for 1 hour at 37°C with shaking at 1,000 RPM. To ensure a single cell suspension, digested organoids cells were forced through a cell strainer (70 μm) into a 50 ml conical tube containing STI. Following cell count and viability determination, the resulting single cell suspension was subjected to organoid formation assays as described for 14 days.

### Mitochondrial Assays

The activity of mitochondria Complex I was measured in peeled murine esophageal epithelium using Complex I Enzyme Activity Assay kit (Abcam, ab109721) according to the manufacturer’s instructions. Briefly, peeled murine esophageal epithelium from young and aged mice was suspended in 500 µl chilled PBS and completely homogenized using a Dounce homogenizer with 20-40 passes. Protein lysis buffer was added to the tissue for protein extraction followed by 30 min ice incubation to allow solubilization. Samples were centrifuged at 16,000 x g for 20 min at 4° C. Supernatant was collected as tissue lysate and diluted to a desired concentration after protein estimation. Tissue lysate was added to 96-well microplates precoated with capture antibodies specific for Complex I. Once target was immobilized, Complex I activity was determined following the oxidation of NADH to NAD and the simultaneous reduction of a dye. Absorbance was measured at OD=450 nm using a spectrophotometer. mtDNA level was measured by qPCR of DNA from peeled murine esophageal epithelium. DNA isolation was performed using DNeasy Blood and Tissue Kit (Qiagen Cat# 69506) according to the manufacturer’s instructions. qPCR was performed using PowerUp SYBR green master mix (Thermo Fisher) with the following primers: Ikbβ For: GCTGGTGTCTGGGGTACAGT Rev: ATCCTTGGGGAGGCATCTAC, and mtDNA D-Loop Fwd:ACTATCCCCTTCCCCATTTG Rev: TGTTGGTCATGGGCTGATTA. The relative fold change between samples of mtDNA D-loop was calculated with normalization to the nuclear encoded Ikbβ.

### Statistics

Descriptive statistics are presented as mean ± standard deviation or median (minimum-maximum) for continuous variables and frequency counts (percentages) for categorical variables. Two-sample t-test or Wilcoxon rank-sum test and one-way analysis of variance or Kruskal-Wallis test comparing two and three groups, respectively for continuous variables and chi-square test or Fisher’s exact test for categorical variables were used.

## Data & Code Availability

The authors declare that all data supporting the findings of this study are available within the article and its supplementary information files or from the corresponding author upon reasonable request. All scRNA-seq data files along with their associated metadata have been deposited into the GEO database (GSE165880). Custom scripts are available at (https://github.com/alkarami/Whelan_scRNA_Esophagus_May20).

## Acknowledgements

We thank Jean-Pierre Issa, MD and members of the Fels Cancer Institute for Personalized Medicine for conceptual support. We also thank Yin Fei Tan, PhD and members of the Fox Chase Cancer Center Genomics Facility for technical support. The following NIH grants supported this work: R01DK121159 (KAW).

