## Supplementary Figures for "Single cell transcriptomic analysis reveals cellular diversity of murine esophageal epithelium and age-associated mitochondrial dysfunction"

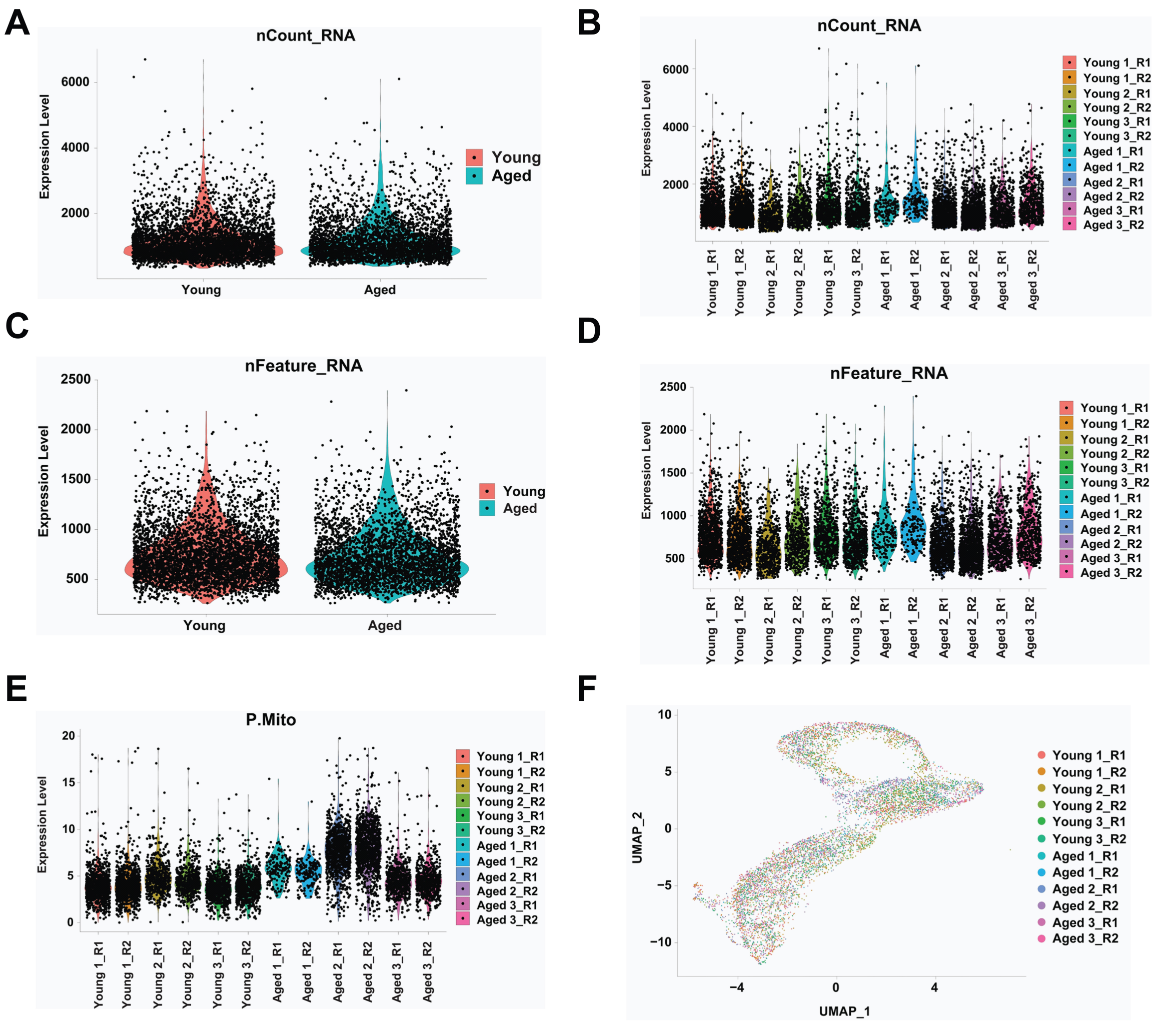


G

| | Young | | | Aged | | |
|---|---|---|---|---|---|---|
| | 1 | 2 | 3 | 1 | 2 | 3 |
| Replicate 1 | 12180763 | 10472916 | 12865327 | 20744792 | 15286566 | 10251195 |
| Replicate 2 | 10902095 | 12496674 | 11419232 | 19587099 | 18715461 | 13162424 |
| Total | 23082858 | 22969590 | 24284559 | 40331891 | 34002027 | 23413619 |

**Supplementary Figure S1. Quality control metrics for scRNA-Seq data**. Young (n=3) and aged mice (n=3) were subjected to droplet-based 3’ scRNA-Seq. 2 reads were done for each mouse, resulting in a total of 12 samples in the analysis. (**A,B**) Transcript counts for young and aged mice are shown by cell in each experimental group (**A**) or in each replicate (**B**). (**C,D**) Unique transcript counts for young and aged mice are shown by cell in each experimental group (**C**) or in each replicate (**D**). (**E**) Percent expression of mitochondrial genes for each cell in each replicate. (**F**) Each cell is colored according to the replicate to which it belongs and the distribution across the UMAP object is shown. (**G**) Number of reads per replicate is shown as along with total read number per sample. *R1, replicate 1; R2, replicate 2.

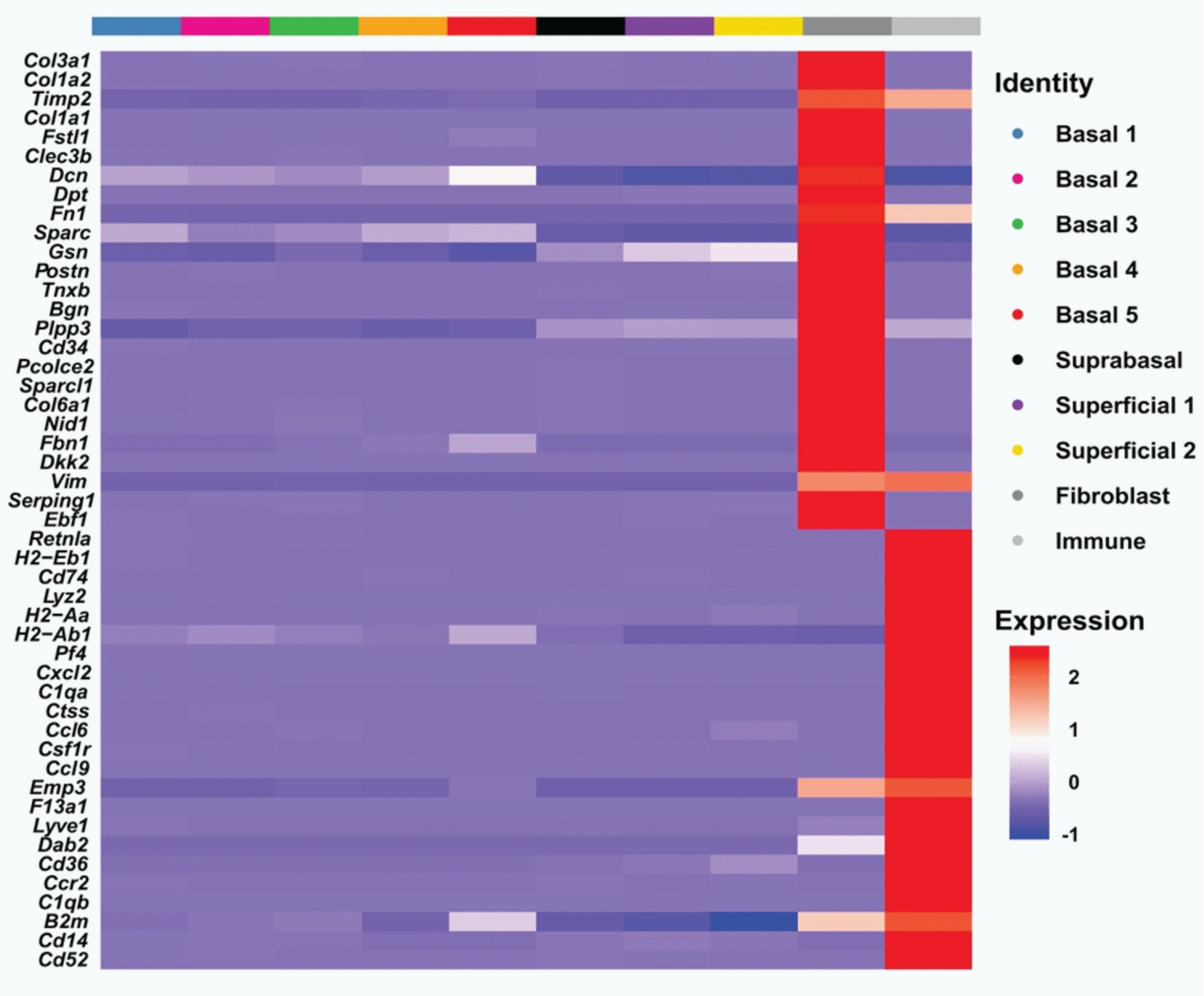


**Supplementary Figure S2. Gene signatures of clusters identified as fibroblast and immune cells in peeled murine esophageal epithelium.** Top 25 upregulated genes in the fibroblast and immune clusters and their respective expression z-scores across all clusters identified in peeled esophageal epithelium are shown.

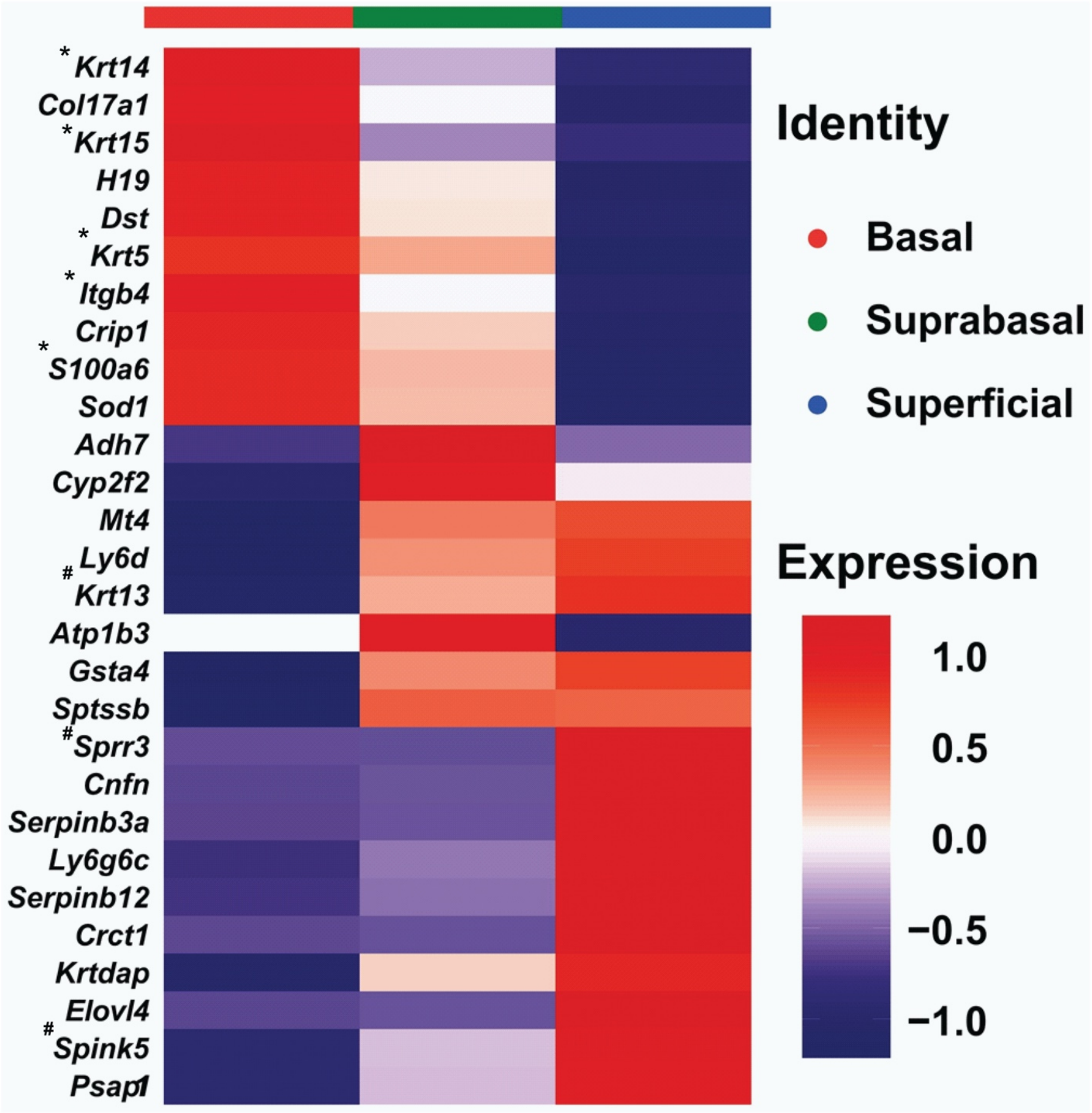


**Supplementary Figure S3. Identification of potential marker genes for basal, suprabasal and superficial subsets in murine esophageal epithelium.** Top marker genes in basal, suprabasal and superficial cells and their respective expression z-scores by cell type. *, indicates known markers of basal esophageal keratinocytes. #, indicates known markers of differentiation in esophageal keratinocytes.

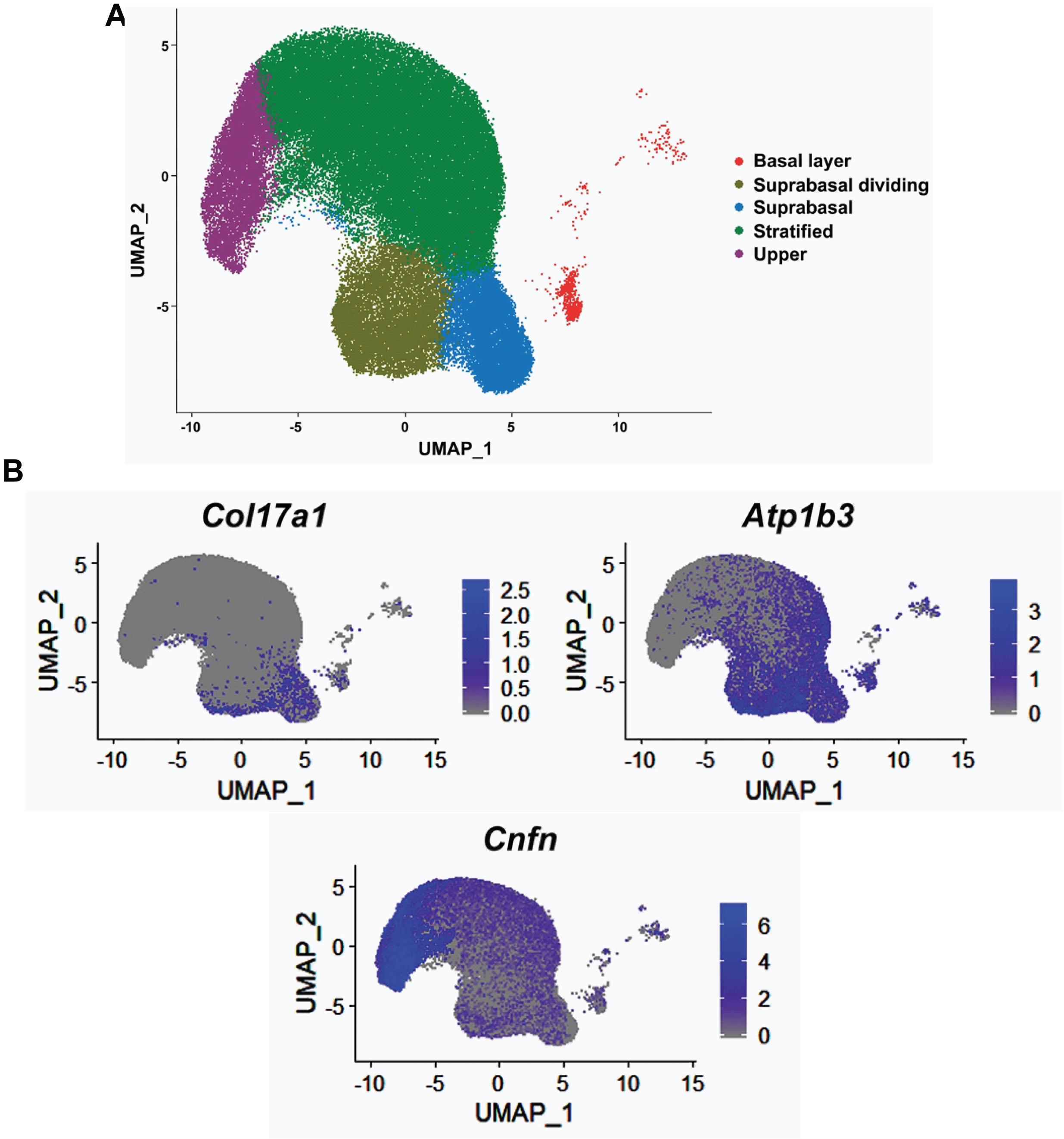


**Supplementary Figure S4. Cellular populations of the human esophageal epithelium.** Single cell RNA-sequencing data was collected from the Tissue Stability Atlas (https://www.tissuestabilitycellatlas.org/oesophagus) and analyzed to identify the cell clusters and localizations of cell type markers in human esophageal epithelium. **(A)** The UMAP plot of annotated cell clusters of the frozen human esophageal epithelium. **(B)** Localization of the novel murine markers for basal (*Col17a1*), suprabasal (*Atp1b3*), and superficial (*Cnfn*) layers in the human frozen esophageal epithelium.

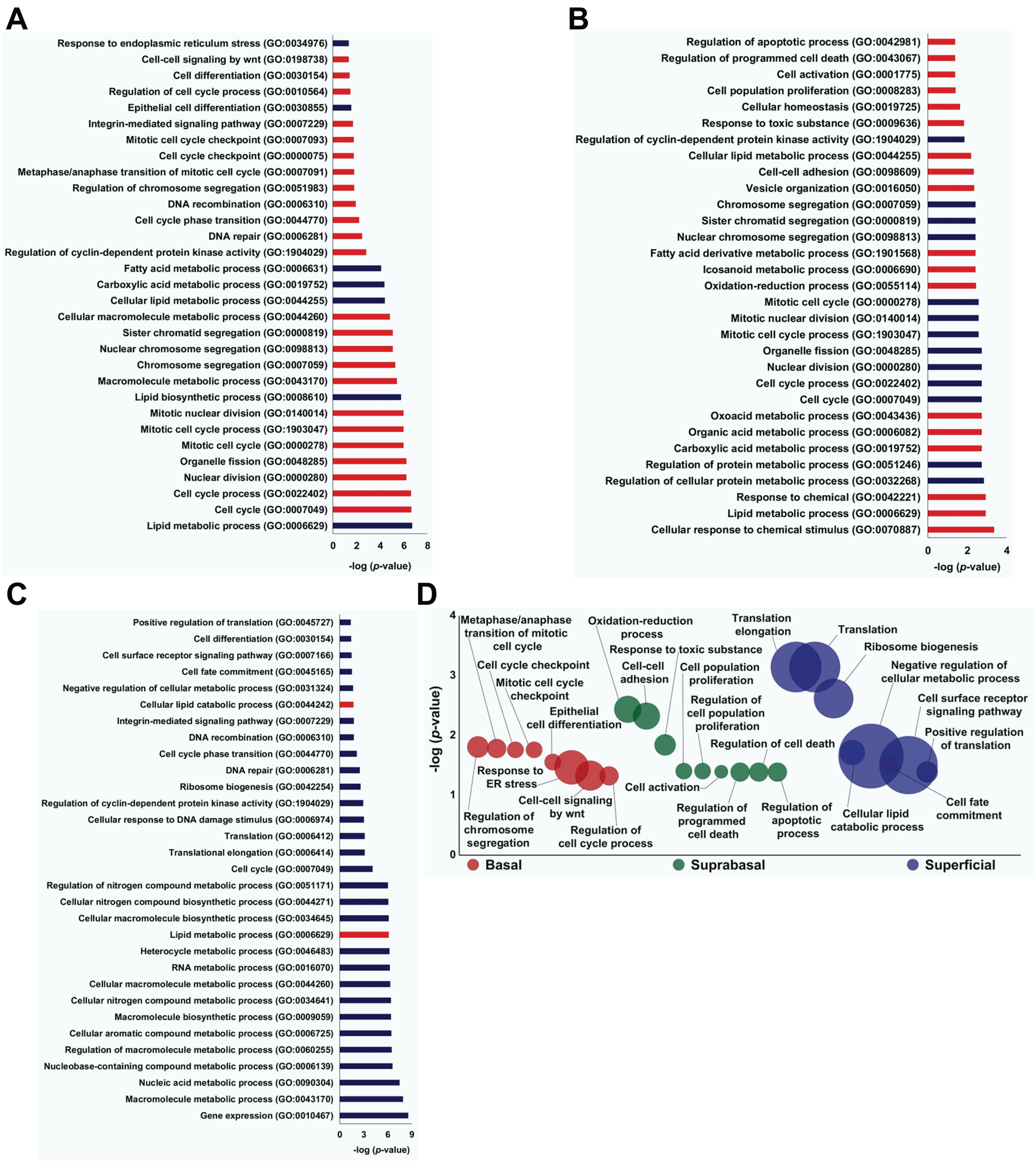


**Supplementary Figure S5. Top biological processes altered in basal, suprabasal and superficial murine esophageal epithelium.** PANTHER analysis categorized the gene ontology (GO): biological processes significantly enriched in basal, suprabasal, and superficial epithelium. (**A-C**) The top GO terms according to *p* value are shown for basal (**A**), suprabasal (**B**) and superficial (**C**) cells, with red denoting over-representation and blue denoting under-representation of indicated biological processes. (**D**) Uniquely enriched biological processes in basal, suprabasal and superficial cells are shown. Circle size reflects number of genes associated with respective biological processes.

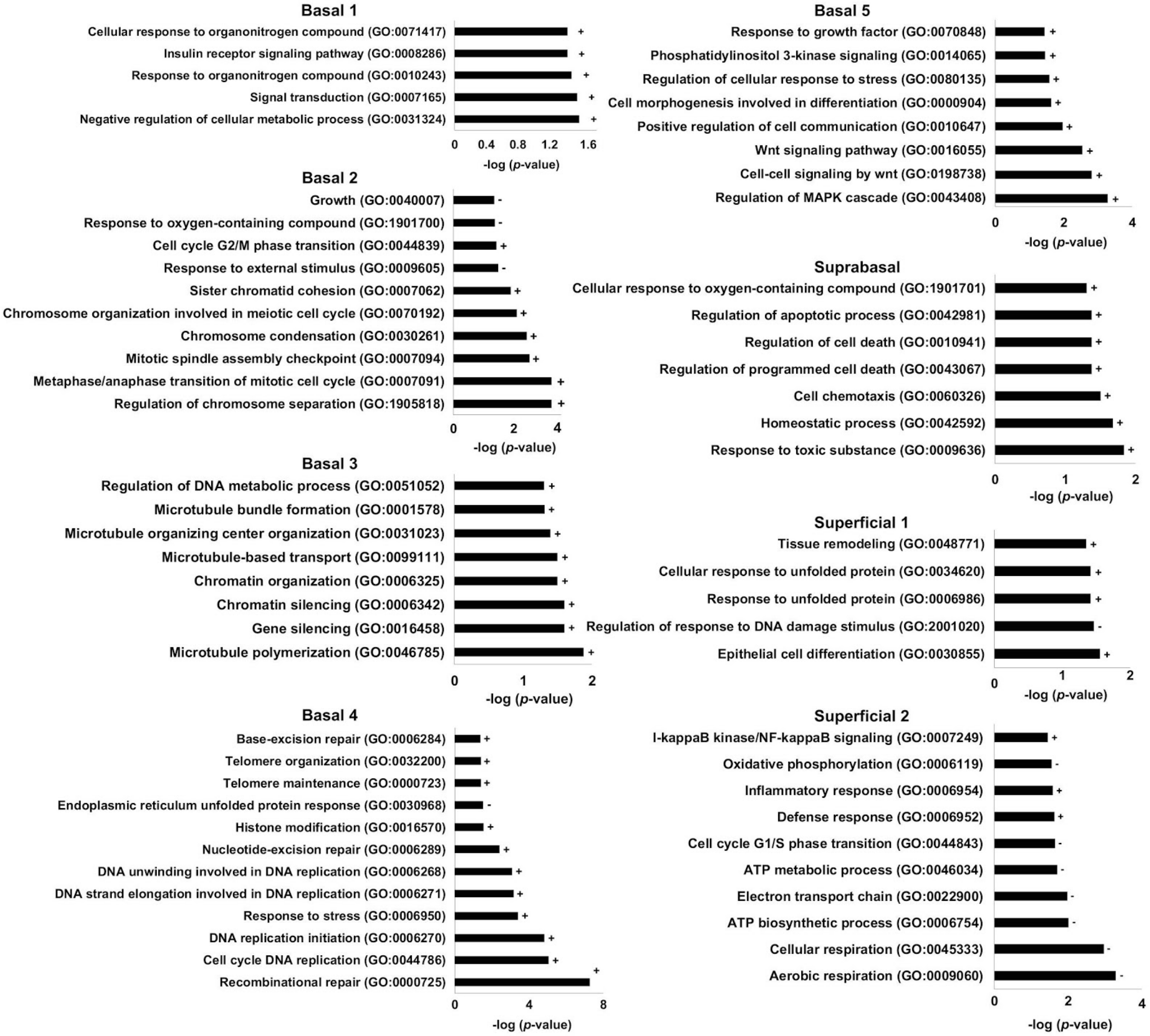


**Supplementary Figure S6. Biological processes altered in individual cell clusters of murine esophageal epithelium.** PANTHER analysis categorized the gene ontology (GO): biological processes significantly enriched in identified cell clusters in murine esophageal epithelium. The unique GO terms are shown with (+) denoting over-representation and (-) denoting under-representation of indicated biological processes.

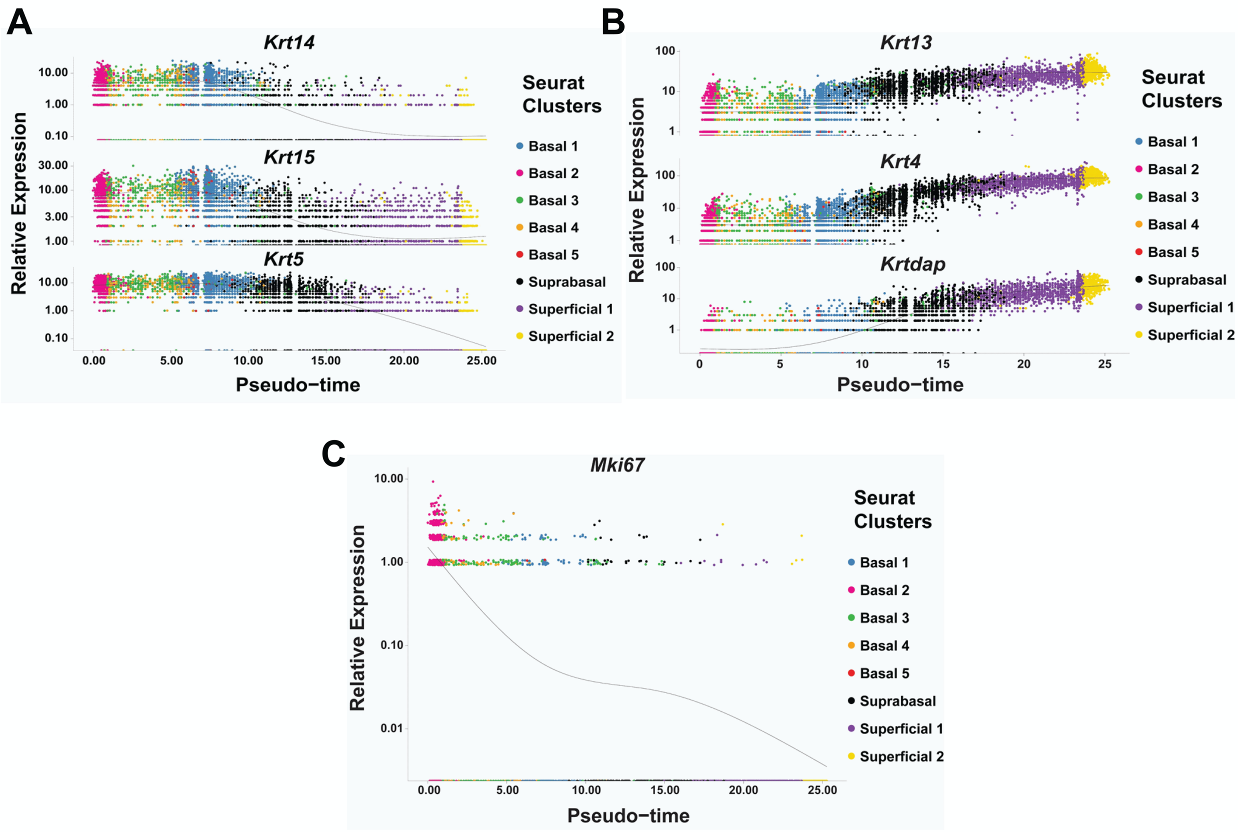


**Supplementary Figure S7. Relative expression of markers of basal cells, differentiation and proliferation along the pseudotime trajectory of the basal-superficial axis**. The relative expression values of basal markers *Krt5*, *Krt14* and *Krt15* (**A**), differentiation markers *Krtdap*, *Krt13* and *Krt4* (**B**), and proliferation marker *Mki67* (**C**) are plotted against pseudo-temporal values that identify basal cluster 2 as the start of the trajectory and superficial cluster 2 as the end.

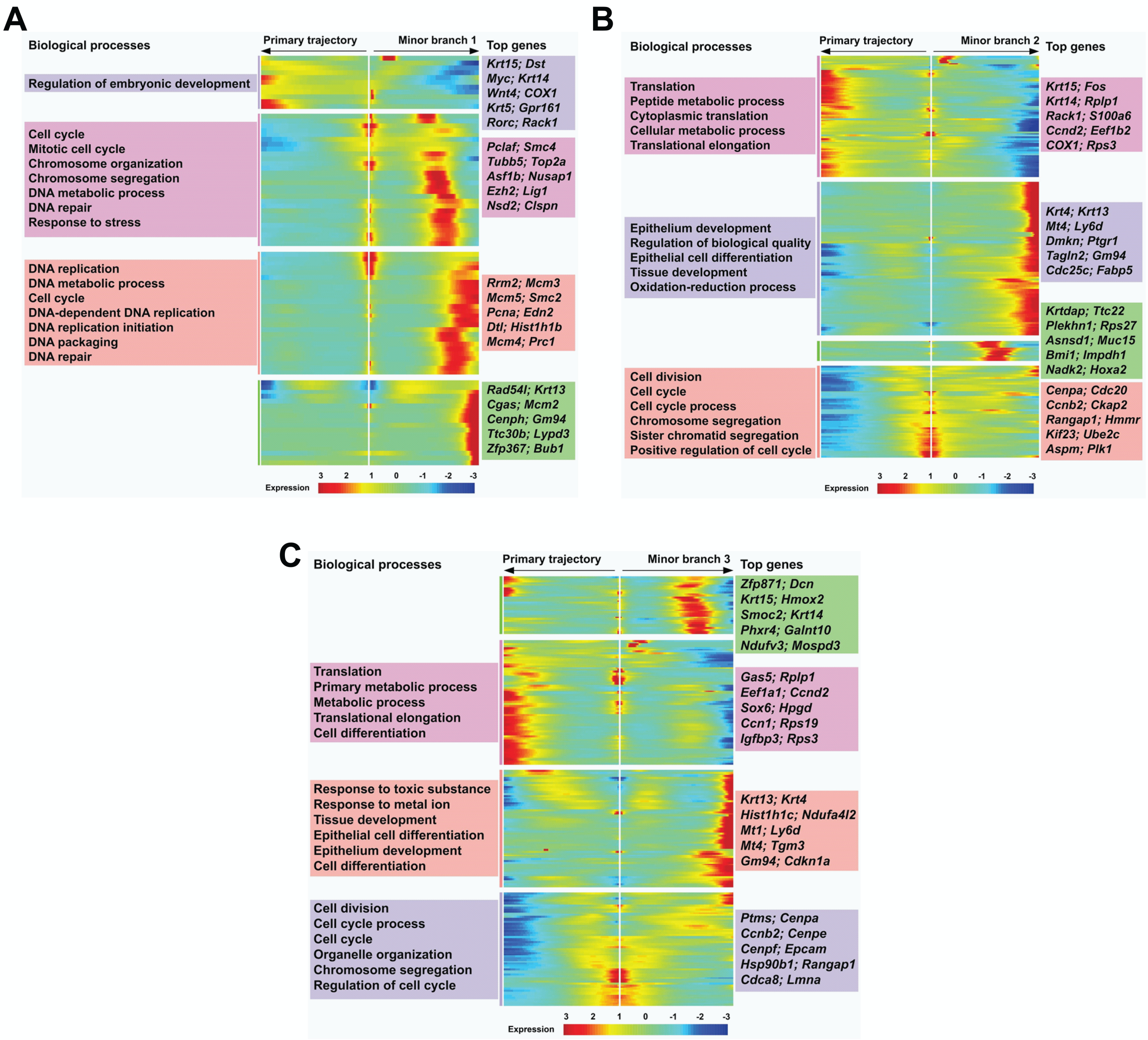


**Supplementary Figure S8. Molecular characterization of minor branches emerging from pseudotemporal analysis of esophageal basal cells**. (**A-C**) Branch-dependent analysis of branch points 1 (**A**), 2 (**B**), and 3 (**C**). Heatmaps represent expression z-scores of genes along the trajectory. STRING analysis revealed biological processes significantly altered in gene clusters. Representative biological processes and top genes are shown.

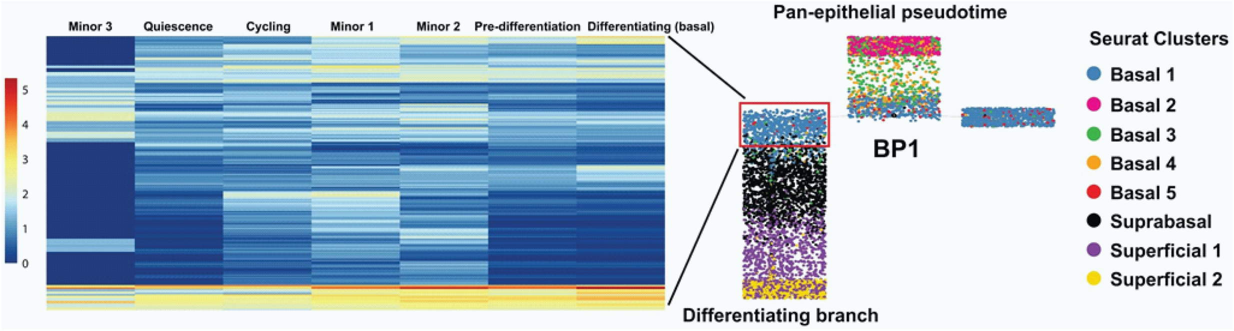


**Supplementary Figure S9. Identification of pre-differentiation branch in esophageal basal pseudotemporal trajectory**. Normalized gene expression profile comparison of individual branches of basal cell pseudotemporal trajectory and basal cells from the differentiating branch of the pan-epithelial pseudotime.

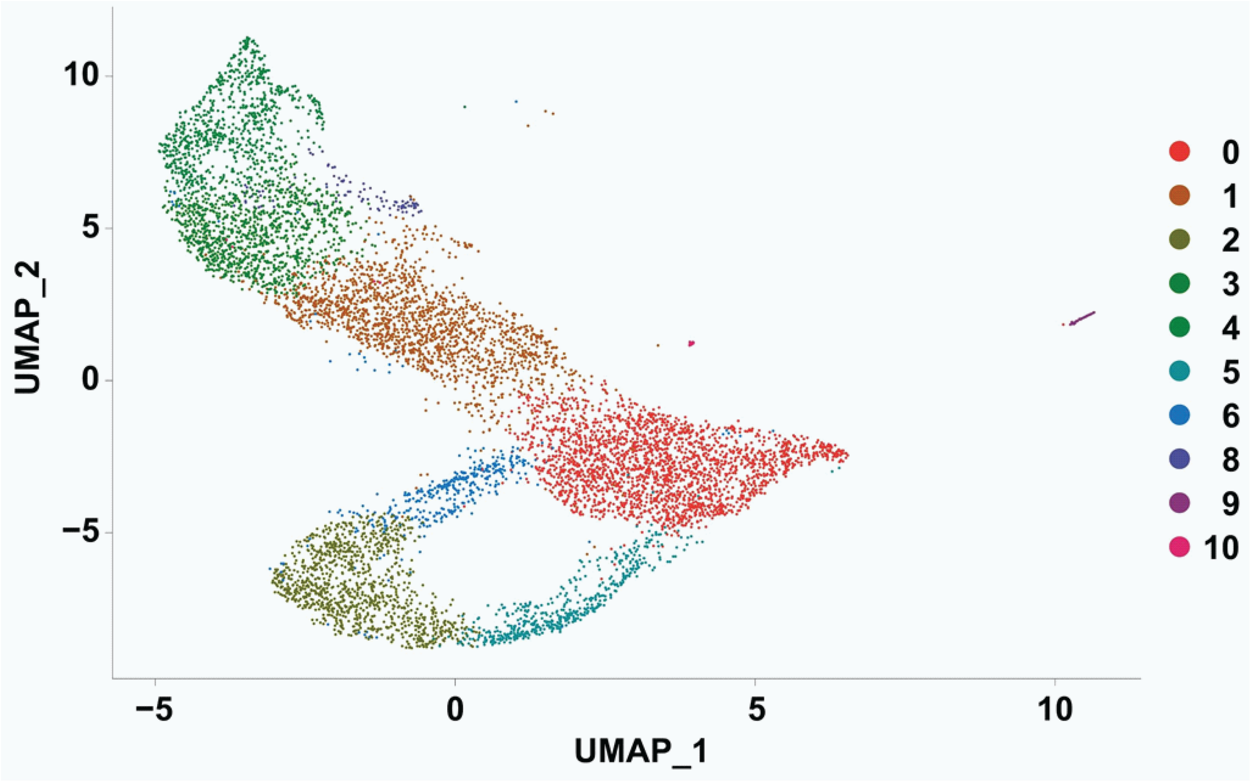


**Supplementary Figure S10. Identification of cell cycle-independent clusters.** Seurat's Uniform Manifold Approximation and Projection (UMAP) was used to identify distinct cell clusters within the epithelial dataset after removal of cell cycle genes. Despite the removal of cell cycle genes in cluster determination, the general formation of the clusters and the cycle formation of clusters are similar to the results of clustering with cell cycle genes included.

**Supplementary Figure**

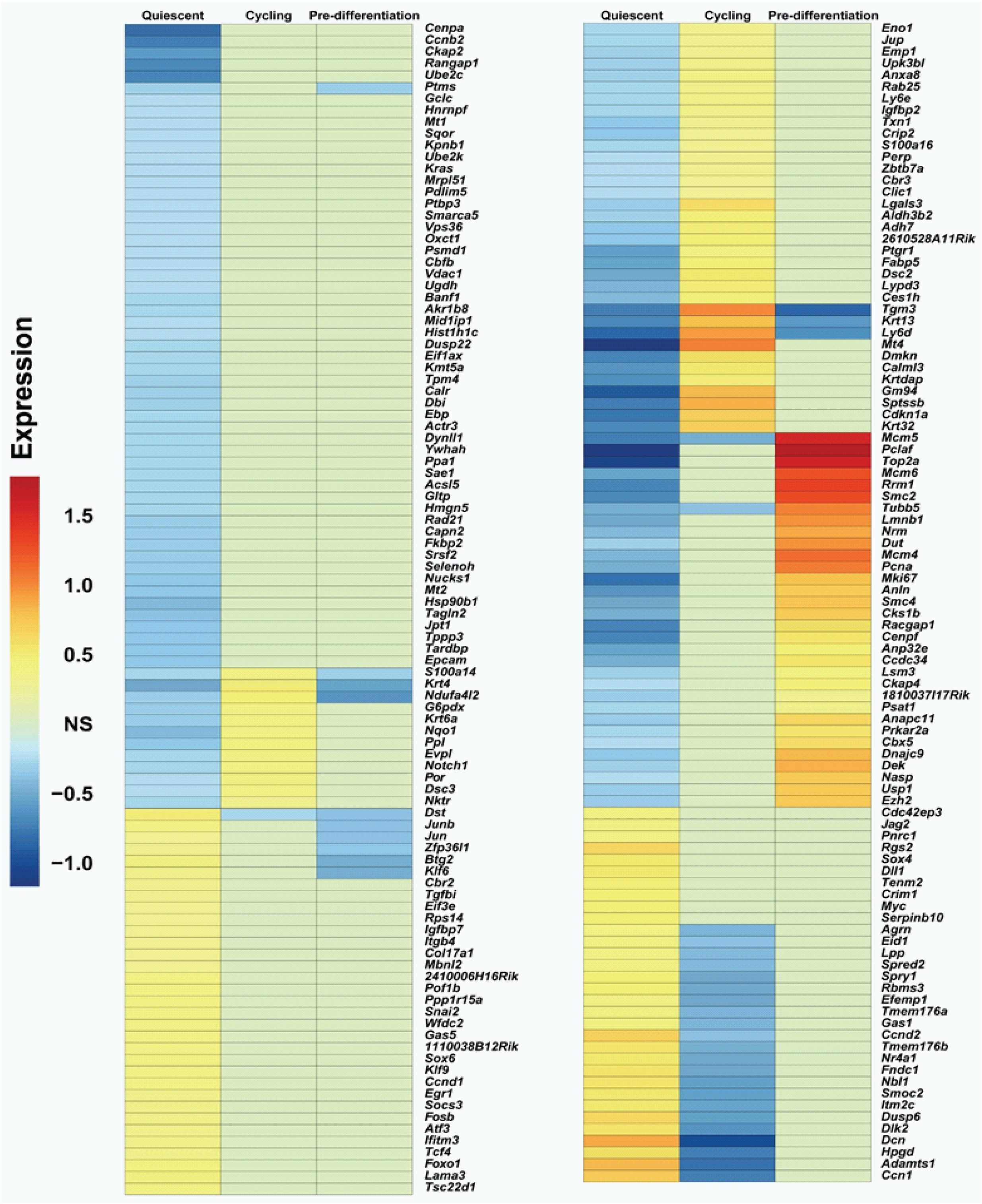


**S11. Characterization of quiescent branch in pseudotemporal trajectory of esophageal basal cells**. Expression fold change comparisons among quiescent, cycling and pre-differentiation branches. NS, non-significant.

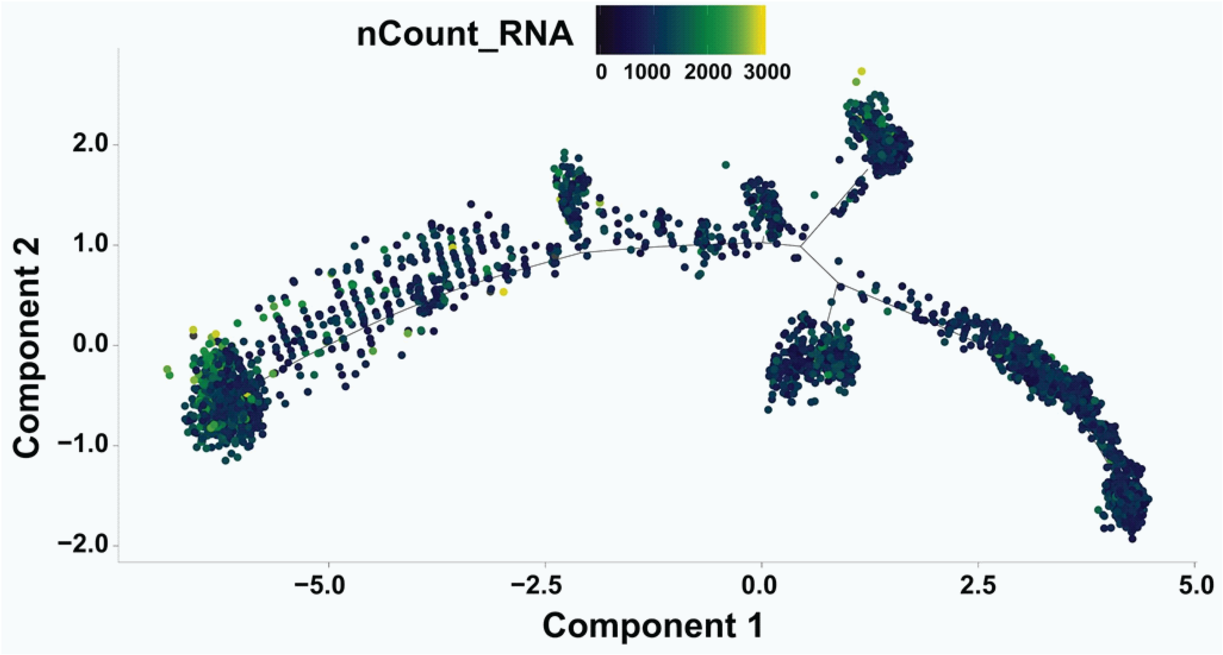


**Supplementary Figure S12. RNA content along the in pseudotemporal trajectory of esophageal basal cells**. Total RNA counts for each cell in the in basal pseudotime trajectory.

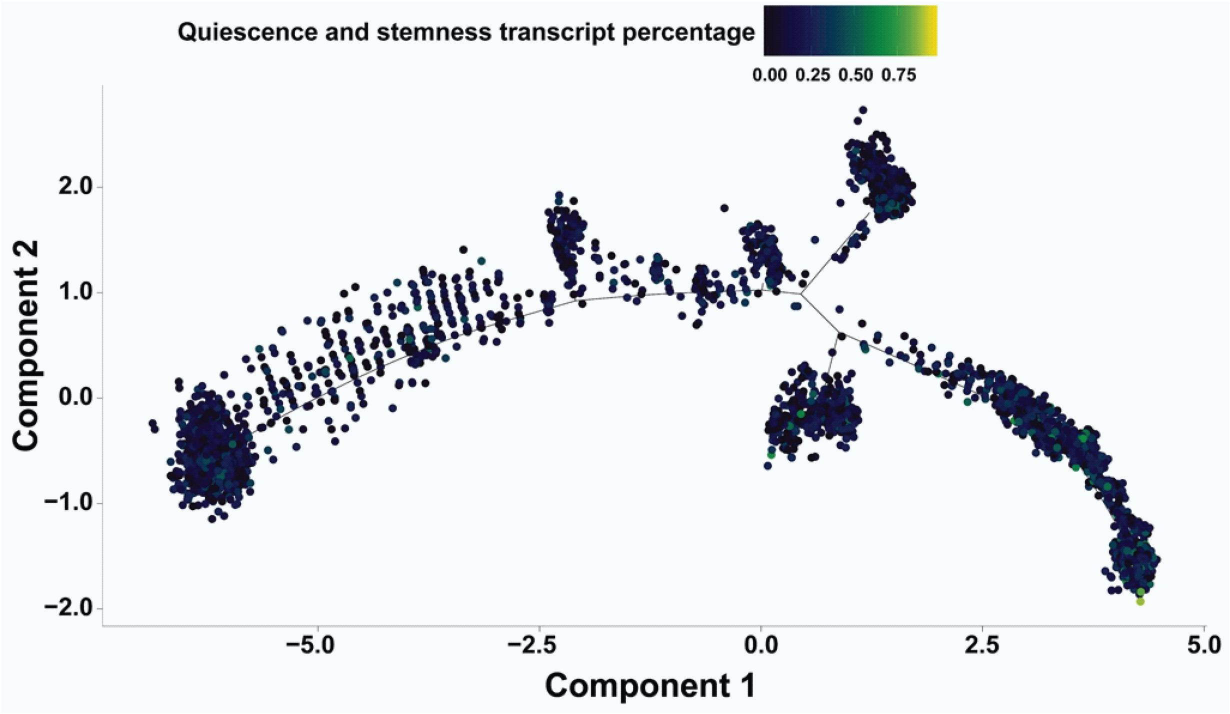


**Supplementary Figure S13. Quiescence and stemness transcripts along the pseudotemporal trajectory of esophageal basal cells**. The percentage of transcripts expressing quiescence and stemness genes as described by Cheung and Rando (2013).

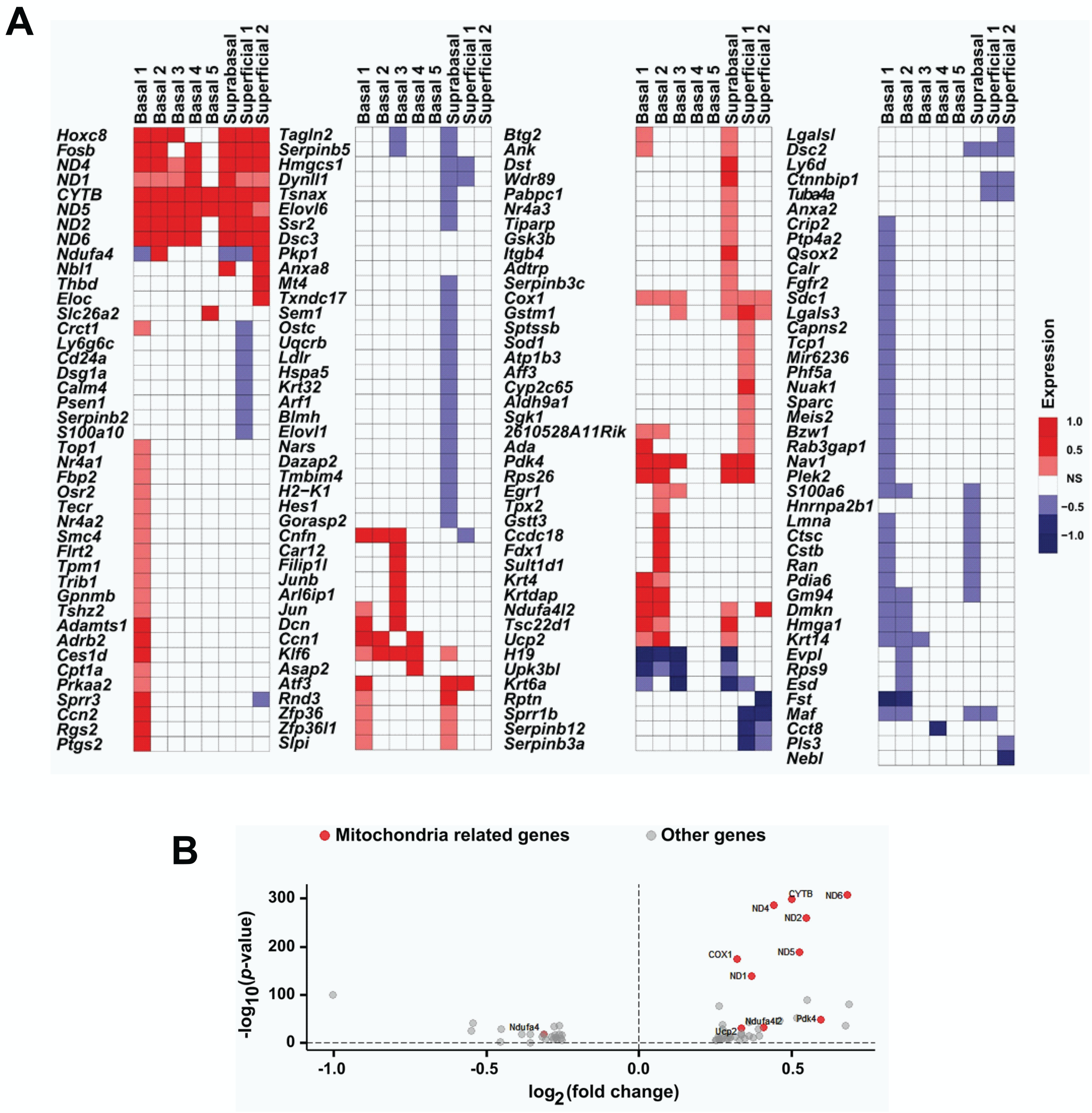


**Supplementary Figure S14. Age-associated alterations in esophageal epithelial gene expression.** (**A**) Heatmap of the fold changes for cluster-specific differentially expressed genes in aged esophageal epithelium. (**B**) Volcano plot of the differentially expressed genes in aged epithelial cells compared to young epithelial cells.
